# *Sox15* is a novel retinal developmental gene that promotes cone photoreceptor differentiation through inhibition of alternative rod photoreceptor fates

**DOI:** 10.1101/2024.12.10.627788

**Authors:** Justin J. Belair-Hickey, Saeed Khalili, Brenda L.K. Coles, Brian G. Ballios, Jeff C. Liu, Gary D. Bader, Derek van der Kooy

## Abstract

The type of cell fate decisions a progenitor can make in the developing nervous system are dependent on a combination of cell intrinsic gene expression programs and cell extrinsic signaling molecules that influence expression programs. Moreover, both the progenitor’s spatial location in the tissue and time at which it exists during development can, to varying degrees, influence the competency of differentiation. Here, we use the developing mouse retina to understand how signaling factors in the retinal niche can influence a progenitor to become unipotentially-restricted to differentiate into cone photoreceptors, and how this competency changes over the course of retinal development. We show that the secreted inhibitory protein COCO causes progenitors to become cone-restricted, and that the competency for this restricted differentiation is maintained throughout retinal development. Using RNA-sequencing, we identify the transcription factor *Sox15* as a potential novel marker of restricted progenitors in the developing retina, and that it acts in response to COCO’s effects. Knockdown experiments indicate that this novel retinal transcription factor promotes cone differentiation by inhibiting differentiation into alternative photoreceptor types.

## Introduction

Cellular diversity in the central nervous system is generated through a limited number of proliferating progenitor cells with varying degrees of cell fate (differentiation) potential (Molyneaux et al., 2007; Dessaud et al., 2008; Zhang et al., 2023). Within an anatomically defined region of the nervous system, the differentiation potential of any individual progenitor can fluctuate widely, with some being highly multipotential and others restricted to producing one or two cell types (Turner et al., 1990; Franco et al., 2012; Emerson et al., 2013; Gao et al., 2014; Khalili et al., 2018). In addition, there are competing models suggesting that progenitor differentiation can be explained by either pre-determined bias or temporal shifts in competence within individual progenitors (Desai and McConnell, 2000; Lodato and Arlotta, 2015). Studies in molecular developmental biology have shown that this can be influenced by varying combinations of cell intrinsic gene expression programs and extrinsic factors in the progenitor niche (Dessaud et al., 2008; Cepko, 2014; Zhang et al., 2023). For many years, studying the developing retina has been productive in modelling the cellular and molecular mechanisms by which diversity and patterning are generated in the central nervous system. This is due to its ease of access and manipulation as a tissue, defined and uniform laminar architecture, and stereotyped birth order of a large diversity of resident cell types (Bassett and Wallace, 2012; Cepko, 2014; Zhang et al., 2023).

Classic birth dating studies in the developing rodent retina showed that different retinal cell types are produced in overlapping but distinct temporal peaks (Young, 1985; Rapaport et al., 2004). Subsequent viral lineage tracing suggested that this ordered progression of retinal cell type differentiation is created through highly variable and multipotential retinal progenitor cells (RPCs) that pass-through changing competence states to generate a subset of retinal cell types at any given time (Turner and Cepko, 1987; Holt et al., 1988; Wetts and Fraser, 1988; Turner et al., 1990). However, more recent investigations have demonstrated that there exist consistently definable RPCs that are restricted in their differentiation to give rise to two or a single cell type (Rompani and Cepko, 2008; Hafler et al., 2012; Emerson et al., 2013; Suzuki et al., 2013; Khalili et al., 2018). Various mechanisms have been proposed to explain this RPC cell fate restriction, and a key question that remains is whether this progenitor restriction is due to pre-determined cell intrinsic programs or secreted factors in the retinal environment (Cayouette et al., 2006; Bassett and Wallace, 2012; Cepko, 2014; Zhang et al., 2023). Moreover, there are some models that propose a molecularly stochastic influence on RPC differentiation potential (Gomes et al., 2011; He et al., 2012).

In this study, we explore the extent to which cell extrinsic and intrinsic factors affect the ability of an RPC to become restricted to differentiating into the cone photoreceptor cells of the mammalian retina, and how cone differentiation competency changes over the time course of retinal development. This specific retinal lineage is an important area of investigation due to the existence of high acuity, cone-dense regions in some vertebrate retinas (Bringmann et al., 2018; Choi et al., 2024). From a cell fate perspective, these cone-dense areas of the retina represent a dramatic reduction in cellular diversity compared to the surrounding retinal tissue, and proposed explanations for this cell type restriction include selective cell death, migration, or the existence of unipotential RPCs (Diaz Araya and Provis, 1992; da Silva and Cepko, 2017; Bringmann et al., 2018; Choi et al., 2024). In addition, understanding the mechanisms by which RPCs can become fate-restricted to cone photoreceptors is translationally important for efforts to replace human cone photoreceptors lost due to disease or damage. Blinding eye diseases, such as macular degeneration, involve a selective loss of cones resulting in severe reduction in high acuity and central vision (Fritsche et al., 2014).

Compared to cell-intrinsic gene regulatory programs, much less is known about how signaling factors in the retinal niche can influence photoreceptor differentiation from RPCs (Livesey and Cepko, 2001; Swaroop et al., 2010; Cepko, 2014; Brzezinski and Reh, 2015). Furthermore, niche factors that can drastically restrict RPCs to unipotential differentiation are seldom identified. We chose to focus on a retinal niche factor called COCO, which is a multifactorial inhibitor of TGFβ, WNT, and BMP signaling (Bell et al., 2003; Zhou et al., 2015). COCO is a member of a group of secreted inhibitory proteins important for neural induction through a default mechanism (Ozair et al., 2013). Accordingly, COCO has been shown to cause a biased or restricted differentiation in RPCs to cones, which is thought to represent a default mechanism of photoreceptor differentiation in the retina (Adler and Hatlee, 1989; Mears et al., 2001; Swaroop et al., 2010; Zhou et al., 2015; Khalili et al., 2018). Our data presented here suggest that exposure to COCO can cause RPCs to generate only cone photoreceptors in mouse and human developmental systems and in multiple cell culture paradigms. RPCs also seem to maintain the competency to be cone-restricted when exposed to COCO’s inhibitory influence regardless of their embryonic age. We hypothesized that there should be a molecular signature of cone-restricted progenitors in response to COCO’s effects. Using RNA-sequencing, we identified a transcription factor, *Sox15*, previously unknown to be involved in any aspect of vertebrate retinal biology. We propose that *Sox15* may be such a molecular signifier of an RPC that is committed to differentiate into cone photoreceptors, and that it functions to promote cone fate through inhibiting differentiation to alternative photoreceptor cell types. Given the highly conserved molecular mechanisms of retinal differentiation across vertebrate and invertebrate species (Swaroop et al., 2010; Gehring, 2014; Zhang et al., 2023), we hypothesize that secreted proteins in the retinal niche inhibit RPCs from differentiating along alternative pathways, thereby promoting cone specification by default. This may be a shared strategy for modulating cone density and distribution in the retina in many vertebrate species.

## Results

To probe how secreted retinal niche factors affect RPC differentiation, we chose to use a colony forming sphere assay. In this assay, single RPCs in suspension culture will proliferate to give rise to a clonally derived free-floating sphere colony (Tropepe et al., 2000; Coles-Takabe et al., 2008). This is an ideal culture system that allows one to select for single proliferative progenitors at different timepoints in a tissue where there is a mixture of post-mitotic differentiating cells (from the dissection) that may obscure interpretations of differentiation potential. It avoids also the need for FACS or MACS techniques that may cause excessive amounts of cell death or biased survival of certain cell types. Unlike colonies derived from stem cells, progenitor colonies have limited self-renewal and are unable to be serially passaged (Barrandon and Green, 1987; Balenci and van der Kooy, 2014). Once colonies have formed, individual suspension colonies are easily picked and plated at a single colony per well in various tightly controlled differentiation conditions. The resulting differentiated cells then represent a clone of progeny originating from a single progenitor cell.

### Cone photoreceptor restricted differentiation potential is maintained in RPCs throughout embryonic retinal development

Single RPC colonies isolated from various murine retinal embryonic ages were differentiated for 4 weeks in pan-retinal (PAN) or COCO conditions. PAN is a minimal medium (see Methods) that causes undirected multipotential differentiation to all major retinal cell types (Tropepe et al., 2000; Coles et al., 2004), and our group and others have shown previously that COCO is present in the developing neural retina and can cause RPCs to become cone differentiation-restricted (Zhou et al., 2015; Khalili et al., 2018). RPCs were derived from embryonic day 12 and 14 (E12, E14); these are time points when high levels of cone differentiation are occurring *in vivo*. We also derived RPCs from E19, which is a late embryonic time point when there is very little to no cone genesis occurring (Young, 1985; Rapaport et al., 2004; Bassett and Wallace, 2012). Under PAN conditions, all time points showed moderate cone photoreceptor differentiation through protein expression of cone arrestin (ARR3) and s-opsin (OPN1SW), the first of which is found in all cones and the second that is highly specific to short-wavelength cones (s-cones). The percentage of all cells in each clone expressing cone markers was not significantly different across embryonic time points (E12 ARR3: 38.91% ± 11.40; E14 ARR3: 30.97 ± 3.79; E19 ARR3: 42.74% ± 5.36; E12 OPN1SW: 28.31% ± 5.14; E14 OPN1SW: 49.86% ± 5.67; E19 OPN1SW: 47.19% ± 5.39; Fig. 1A-C). In contrast, when exposed to COCO, nearly all cells in each clone from every embryonic age were expressing cone markers (E12 ARR3: 82.82% ± 1.95; E14 ARR3: 80.80% ± 4.37; E19 ARR3: 71.84% ± 10.95; E12 OPN1SW: 87.92% ± 2.16; E14 OPN1SW: 92.95% ± 1.90; E19 OPN1SW: 94.07% ± 1.35; Fig. 1D-F). Like PAN conditions, there was no significant difference between embryonic ages in cone differentiation level in COCO (Fig. 1E,F). As we are representing these data as a percentage of cells in each colony, one possible interpretation of the cone bias in COCO conditions is selective survival of cone precursors or differentiated cones. However, when looking at overall clone size at the end of the differentiation period, there were no significant differences between PAN and COCO differentiation conditions at each embryonic time (E12 PAN: 2612 cells ± 878.9 versus E12 COCO: 2157 cells ± 710.2; E14 PAN: 249.4 cells ± 83.1 versus E14 COCO: 446.2 cells ± 151.3; E19 PAN: 99.70 cells ± 28.81 versus E19 COCO: 171.2 cells ± 87.04; Fig. 1G-I). There was, on average, larger clone sizes at earlier embryonic stages (from presumably higher RPC proliferative rates), however, this was not affected by PAN versus COCO differentiation condition (Fig. 1G-I). These data also argue against a selective proliferative advantage in COCO as a reason for the large cone numbers in each clone. We wondered as well whether the cone differentiation effects were specific to our clonal sphere colony culture system. Therefore, we differentiated RPCs from dissociated whole embryonic neural retina plated at low density and observed similar enhancement of cells expressing OPN1SW when exposed to COCO (PAN: 45.9% ± 9.67 versus COCO: 88.58% ± 5.42; Fig. 1J,K). Furthermore, we employed a 3D mouse retinal organoid culture system, where RPCs give rise to not only the full complement of retinal cell classes but also more accurate laminar architecture and morphology of retinal cells (Eiraku et al., 2011). When exposed to COCO, retinal organoids did not show any difference in retinal progenitor marker expression (Fig. S1A), but there was a significant upregulation in the general photoreceptor marker *Crx* and the cone-specific opsin (*Opn1sw*) (*Crx*: 22.59-fold ± 4.330, *Opn1sw*: 12.58-fold ± 1.128; Fig. S1B,C). This apparent differentiation effect was specific to cones, as there was no significant upregulation in *Rho* (rod specific) in COCO compared to PAN differentiation conditions (Fig. S1D). These organoid data are consistent with differentiation effects seen in the 2D (monolayer) RPC culture described above. Overall, these results lead us to propose that the default differentiation pathway from RPCs, regardless of embryonic age, is to an s-cone photoreceptor fate, and that this is achieved through inhibition of cell extrinsic signaling factors that direct RPCs to alternative retinal fates. Most surprising, this default differentiation seems also to be maintained in RPCs when taken out of their inhibitory niche at later embryonic timepoints when cones are no longer being made *in vivo*.

**Fig. 1.**
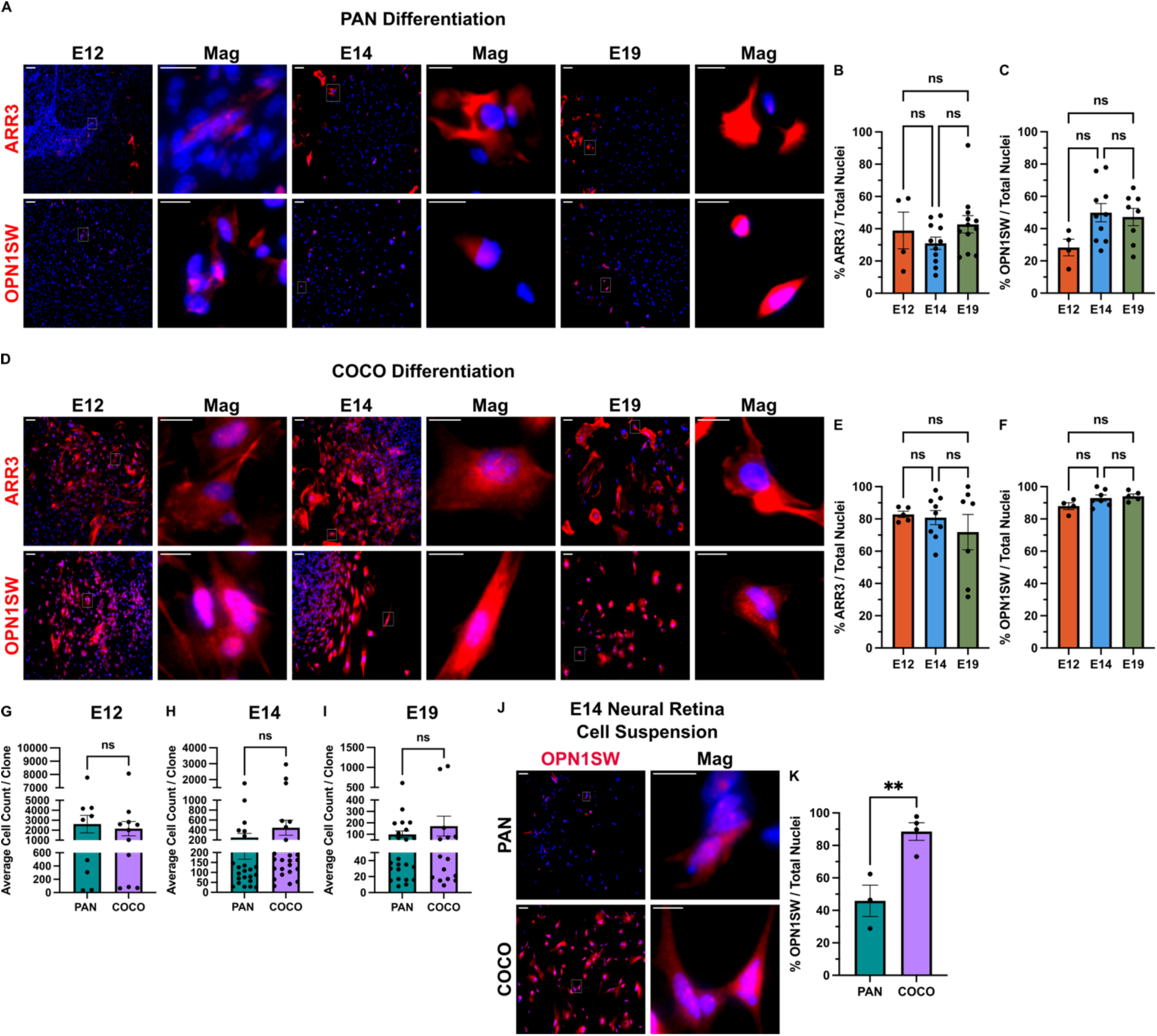
RPCs retain cone differentiation bias in COCO regardless of embryonic age. **(A)** Representative immunofluorescence micrographs of embryonic RPC clones from various ages differentiated in pan retinal conditions (PAN) for 4 weeks. Cells are stained with the cone photoreceptor markers cone arrestin (ARR3) and s-opsin (OPN1SW). **(B)** Quantification of the percentage of totals cells in the clone that are expressing ARR3 (described in **A**). E12 (embryonic day 12), *n* = 4 clones; E14 (embryonic day 14), *n* = 11 clones; E19 (embryonic day 19), *n* = 12 clones; one-way ANOVA with Tukey’s multiple comparisons test, ns= not significant (> *p* = 0.05). **(C)** Quantification of the percentage of totals cells in the clone that are expressing OPN1SW (described in **A**). E12, *n* = 4 clones; E14, *n* = 10 clones; E19, *n* = 8 clones; one-way ANOVA with Tukey’s multiple comparisons test, ns. **(D)** Representative immunofluorescence micrographs of embryonic RPC clones from various ages differentiated in COCO conditions for 4 weeks. **(E)** Quantification of the percentage of totals cells in the clone that are expressing ARR3 (described in **D**). E12, *n* = 5 clones; E14, *n* = 9 clones; E19, *n* = 7 clones; one-way ANOVA with Tukey’s multiple comparisons test, ns. **(F)** Quantification of the percentage of totals cells in the clone that are expressing OPN1SW (described in **D**). E12, *n* = 4 clones; E14, *n* = 7 clones; E19, *n* = 5 clones; one-way ANOVA with Tukey’s multiple comparisons test, ns. **(G-I)** Average number of cells counted (from 4 images) per clone derived from embryonic RPCs differentiated in PAN or COCO conditions for 4 weeks. (**G**) PAN, *n* = 9 clones; COCO, *n* = 11 clones; two-tailed unpaired Student’s t-test, ns, *p* = 0.6885. (**H**) PAN, *n* = 23 clones; COCO, *n* = 24 clones; two-tailed unpaired Student’s t-test, ns, *p* = 0.2659. (**I**) PAN, *n* = 23 clones; COCO, *n* = 15 clones; two-tailed unpaired Student’s t-test, ns, *p* = 0.3680. **(J)** Representative immunofluorescent micrographs of OPN1SW staining in E14 whole neural retinal cell suspensions plated at low density (2 cells / μL) and differentiated in PAN or COCO conditions for 4 weeks. PAN, *n* = 3 clones; COCO, *n* = 4 clones. Error bars represent mean ± SEM. For all micrographs main image scale bar is 100 μm and magnified insert scale bar is 25 μm. **(K)** Quantification of the percentage of totals cells in the clone that are expressing OPN1SW (described in **J**). PAN, *n* = 3 clones; COCO, *n* = 4 clones; two-tailed unpaired Student’s t-test, \*\**p* = 0.0091. Error bars represent mean ± SEM. For all micrographs main image scale bar is 100 μm and magnified insert scale bar is 25 μm.

There are temporal identity transcription factors in the retina that have been shown to influence the competence of RPCs to generate early born versus late born retinal cell types (Elliott et al., 2008; Mattar et al., 2015), and we questioned if COCO may be working to shift the temporal identity of RPCs. We first comparted the expression of temporal identity factors in RPC colonies versus whole neural retina from which they were derived. Regardless of embryonic age of origin, RPC colonies upregulated the late retinal identity factor *Casz1* and downregulate the early retinal identity factor *Ikzf1* (Fig. S2A,B). Given that RPC colonies take 6 days in culture to form, this might reflect an ability of progenitors to “track” developmental time *in vitro* through either an intrinsic counting of cell divisions and/or paracrine or autocrine signaling released within the colony. After 5 days in differentiation culture there was, on average, a greater downregulation of Casz1 in COCO compared to PAN (PAN: - 0.4489-LOG2-fold ± 0.6999 versus COCO: −1.494-LOG2-fold ± 0.6648; Fig. S2C). This indicates that COCO may in part be erasing the late temporal identity in RPCs and that some combination of the WNT, TGFβ, and BMP signaling factors secreted from RPCs is responsible for shifting RPC colonies to a later temporal identity despite being derived from an early embryonic timepoint (E14).

### Cell extrinsic influence on cone restriction in RPCs exists in human retinal development

Given that primate species have a dense, cone photoreceptor-only region in the retina called the fovea (Bringmann et al., 2018), we hypothesized that using COCO to make a cone-restricted progenitor state may be a common mechanism to generate cones among mammals. We utilized two human *in vitro* retinal developmental models to assay the effect of COCO.

In the first, adult retinal stem cells (RSCs) were isolated from human donor eyes. In the ciliary epithelium of the eye, there are quiescent RSCs that, when taken out of their inhibitory niche *in vivo*, can rapidly proliferate to give rise to a colony of cells containing both downstream neural retinal progenitors (RPCs) and retinal pigmented epithelium (RPE) progenitors. These progenitors, which are the progeny of a single RSC, can differentiate into all major cell types in the retina, thereby modelling all retinal developmental lineages (Tropepe et al., 2000; Coles et al., 2004; Khalili et al., 2018). Single clonal RSC sphere colonies can be used, in the same ways as described above, to investigate the effects of cell extrinsic signaling factors on RPC differentiation. After 6 weeks of COCO exposure, there was a large increase in the clonal progeny of human adult RSCs expressing the cone specific OPN1SW when compared to PAN conditions (PAN: 28.10% ± 4.478 versus COCO: 60.58 ± 5.778; Fig. 2A,B). The percentage of cones in each COCO exposed clone closely matched previous data from our lab using mouse RSC colonies, and we have shown that cone number is not as high as from RPC colonies (in Fig. 1) due to the presence of RPE progenitors in the RSC colony. RPE progenitors are developmentally restricted to making RPE *in vivo* and do not normally generate neural retinal cells (Khalili et al., 2018). This photoreceptor differentiation effect was specific to cones, as there was no increase in RHO expression (a rod marker) after COCO exposure (PAN: 0.86% ± 0.5374 versus COCO: 0.19% ± 0.1634; Fig. 2C).

**Fig. 2.**
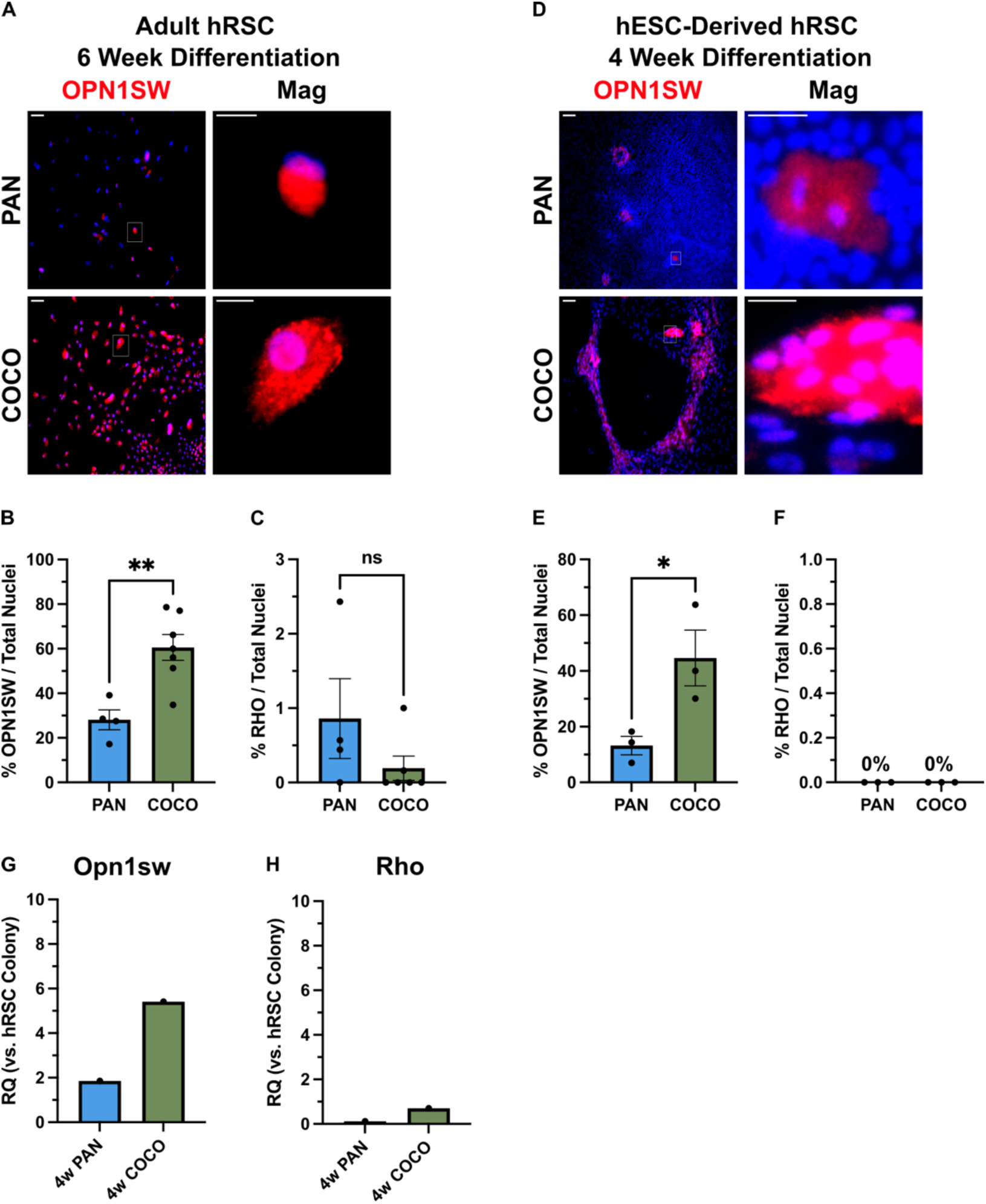
Cone differentiation bias in the presence of COCO is present in progenitors derived from human RSCs. **(A)** Representative immunofluorescent micrographs of clonally derived adult human RSC colonies differentiated in PAN or COCO conditions for 6 weeks. Cells are stained with the cone photoreceptor marker OPN1SW. Quantifications described below are from 4 donor eyes of 2 individuals. **(B)** Quantification of the percentage of totals cells in the clone that are expressing OPN1SW (described in **A**). PAN, *n* = 4 clones; COCO, *n* = 7 clones; two-tailed unpaired Student’s t-test, *\*\*p* = 0.0040 **(C)** Quantification of the percentage of totals cells in the clone that are expressing rod photoreceptor marker rhodopsin (RHO). PAN, *n* = 4 clones; COCO, *n* = 6 clones; two-tailed unpaired Student’s t-test, ns, *p* = 0.1950. **(D)** Representative immunofluorescent micrographs of clonal human RSC colonies derived directly from human embryonic stem cells and differentiated in PAN or COCO conditions for 4 weeks. **(E)** Quantification of the percentage of total cells in the clone that are expressing OPN1SW (described in **D**). PAN, *n* = 3 clones; COCO, *n* = 3 clones; two-tailed unpaired Student’s t-test, *\*p* = 0.0403. **(F)** Quantification of the percentage of totals cells in the clone that are expressing RHO. PAN, *n* = 3 clones; COCO, *n* = 3 clones. **(G-H)** qPCR on 4 week differentiated retinal cells derived from human RSCs. Fold change (RQ) relative to human RSC colonies prior to differentiation. *Gapdh* and *Actb* were used as endogenous control genes. *N* = 1 experiment of cells collected from 2 donor eyes of one individual. Error bars represent mean ± SEM. For all micrographs main image scale bar is 100 μm and magnified scale bar is 25 μm.

This previous assay was done using an adult tissue-specific stem cell, and we questioned whether COCO mediated cone restriction was possible in RPCs that are more representative of early human embryonic retinal development. Human embryonic stem cells (hESCs) can undergo a reliable and spontaneous differentiation towards a retinal fate in minimal medium conditions (Clarke et al., 2012). In this assay, patches of cells will begin to express early eye field transcription factors known to specify which region of the developing brain will give rise to the optic vesicle and later retina (Zuber et al., 2003; Clarke et al., 2012). Within these patches, there is a small number of pigmented RSCs that behave very similar to their adult counterparts (Clarke et al., 2012), and so we used these hESC-derived RSC colonies to test the effect of COCO on human embryonic retinal cell fate specification. As seen with adult RSCs, COCO exposure caused a large increase in OPN1SW expressing cells in each clone when compared to PAN conditions (PAN: 13.21% ± 3.284 versus COCO: 44.65% ± 9.984; Fig. 2D,E) and no increase in RHO expressing cells (PAN: 0% versus COCO: 0%; Fig. 2F). Additionally, qPCR analysis of *Opn1sw* was consistent with what was observed with protein staining (PAN: 1.852-fold versus COCO: 5.411-fold; Fig. 2G); there was also only a very small effect on *Rho* expression (Fig. 2H). These data suggest that COCO mediated inhibition may be a common mechanism in the mammalian retina to differentiate s-cone photoreceptors, leading to our hypothesis that modulating the concentration of COCO in subregions of the retina (such as the fovea) may be a way to generate increased cone density.

### *Sox15* is upregulated in RPCs undergoing cone-restricted differentiation

We hypothesized that there should be a unique molecular signature of RPCs as they are undergoing cone-restricted differentiation in the presence of COCO, and that there may be novel transcription factors that are functionally involved in this process. Starting with murine E14 RPC colonies, we carried out bulk RNA-sequencing (seq) at various time points along cone differentiation in the presence of COCO (Fig. 3A). We analyzed as well a time course of differentiation in the presence of taurine and retinoic acid (T + RA), as our group has shown previously that this combination of retinal niche signaling factors can cause a rod-restricted differentiation in RPCs (Ballios et al., 2012; Khalili et al., 2018). Principle component analysis (PCA) was used to assess how transcriptionally distinct the cultures were at increasing times of differentiation and between the two photoreceptor promoting conditions. Normally RPCs are given 6 days to form a clonal sphere colony in a serum free medium with bFGF and heparin, but we questioned whether priming (supplementing) the colonies with COCO or T + RA during this period would cause an earlier photoreceptor differentiation signature. COCO-primed sphere colonies were very transcriptionally similar to non-primed spheres; however, T + RA priming caused the colony to be more transcriptionally similar to 5 days of T + RA differentiation (Fig. 3B). As predicted, the PCA showed clear transcriptional distinctions between RPCs exposed to COCO versus T + RA. Most surprisingly, these sharp differences were apparent even at 5 days of differentiation when the cells are thought to still be in a proliferative RPC-like state (Fig. 3B). Upon closer inspection of known photoreceptor marker gene expression, COCO-treated cells showed a clear expression of genes specific to fully differentiated cones as time progressed (Fig. 3C). Early differentiation time points did show some expression of known rod photoreceptor genes, however, these were downregulated at later time points (Fig. 3B). In accordance with our protein staining data, the cells did not express medium wavelength cone (m-cone) markers and are, therefore, likely s-cone photoreceptors (Fig 3B). This is also to be expected since differentiation to m-cones has been shown to require thyroid hormone signaling (Ng et al., 2001; Roberts et al., 2006; Eldred et al., 2018), and we do not have thyroid hormone in our tissue culture medium.

**Fig. 3.**
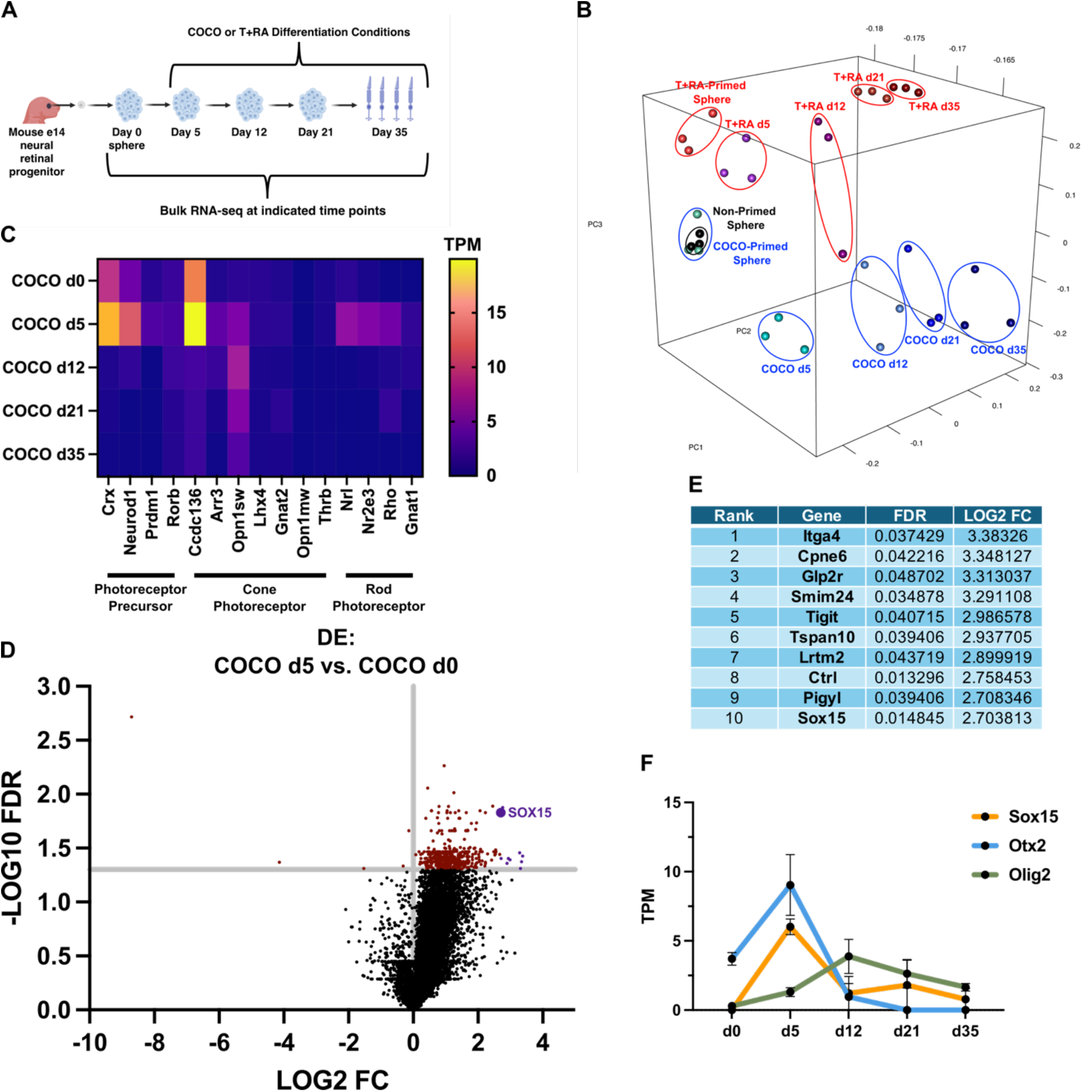
*Sox15* is a novel retinal gene upregulated early in RPCs exposed to cone differentiation conditions. **(A)** Experimental outline for bulk RNA sequencing from E14 RPCs in photoreceptor differentiation conditions. **(B)** 3D principal component analysis of whole transcriptomes from retinal cells at various time points along cone (COCO) or rod (T + RA) photoreceptor differentiation conditions. Primed spheres contain COCO or T + RA in cell culture medium during seven-day colony formation prior to adherent differentiation. Non-primed spheres are grown in baseline SFM without differentiation factors prior to adherent differentiation. Biological replicates are circled in the plot; *n* = 3 biological replicates per timepoint and growth condition. **(C)** Normalized heatmap expression (from sequencing described in **A**) quantified as transcripts per kilobase million (TPM) of photoreceptor precursor, cone, and rod markers across COCO differentiation timepoints. d0 represents COCO primed spheres prior to adherent differentiation. **(D)** Volcano plot of differentially upregulated and downregulated genes in day 5 (d5) COCO cells compared to d0 (COCO-primed sphere) cells. X-axis shows fold change (FC) and Y-axis shows false discovery rate (FDR), with an intersecting line indicating genes above a -LOG10 FDR of 1.3 (0.05). Any gene at or above this is considered to be significantly differentially expressed (red dots). The Ballgown software package was used for differential expression and the top ten differentially upregulated genes are highlighted in purple. Pseudogenes and non-coding RNA were filtered out of the differential expression analyses. **(E)** List of top ten significantly differentially expressed genes (described in **D**). **(F)** TPM across COCO differentiation timepoints (described in **A**) for *Sox15* and two additional genes known to be involved in biasing RPCs towards cone differentiation (*Otx2* and *Olig2*). Error bars represent mean ± SEM.

We predicted that day 5 (d5) of differentiation was when we would identify transcription factors important for a cone-restricted RPC state. In examining the most differentially upregulated genes between COCO d5 and RPC colonies prior to differentiation (d0), there was one transcription factor, *Sox15*, in the top 10 list (Fig. 3D,E). *Sox15* has no known roles in vertebrate retinal development and was previously characterized as a gene important for muscle stem cell repair of damaged tissue (Lee et al., 2004). There are, however, some previous results that made us consider *Sox15* as a promising candidate for a possible role in retinal cell fate specification. In xenopus, the orthologous gene *soxD* is involved in anterior neural induction and its dominant negative form causes impairment in anterior neural development, including an absence of eye formation (Mizuseki et al., 1998). In addition, it is upregulated by chordin, which has similar functional activity to COCO (Deglincerti et al., 2015), and *soxD* is inhibited by BMP4 (Mizuseki et al., 1998). In mouse ESCs *Sox15* is important for neural induction (Choi et al., 2023) and *Sox15* also may have the ability to inhibit Wnt signaling (Moradi et al., 2017). Finally, there are results from a cancer study that *Sox15* is in a gene regulatory network with photoreceptor genes such as *Crx* and *Nrl* (Garancher et al., 2018). When looking at the expression of *Sox15* in our COCO RNA-seq experiment, there was a clear early peak that was rapidly downregulated after day 5, and this followed closely the expression of *Otx2* and *Olig2* (*Sox15* d0: 0 TPM; *Sox15* d5: 6.018 TPM ± 0.564; *Sox15* d12: 1.218 TPM ± 1.218; *Sox15* d21: 1.805 TPM ± 1.805; *Sox15* d35: 0.785 TPM ± 0.785; Fig. 3F), two genes known to be important for biasing cone photoreceptor differentiation (Hafler et al., 2012; Emerson et al., 2013). Most d5 COCO cells had SOX15 expressed at the protein level (85.37% ± 3.507; Fig 4.A,B), and as predicted, this is a stage where there are still proliferating RPCs that do not express mature cone markers like ARR3 (KI67: 35.90% ± 4.891; Fig. 4C-F).

**Fig. 4.**
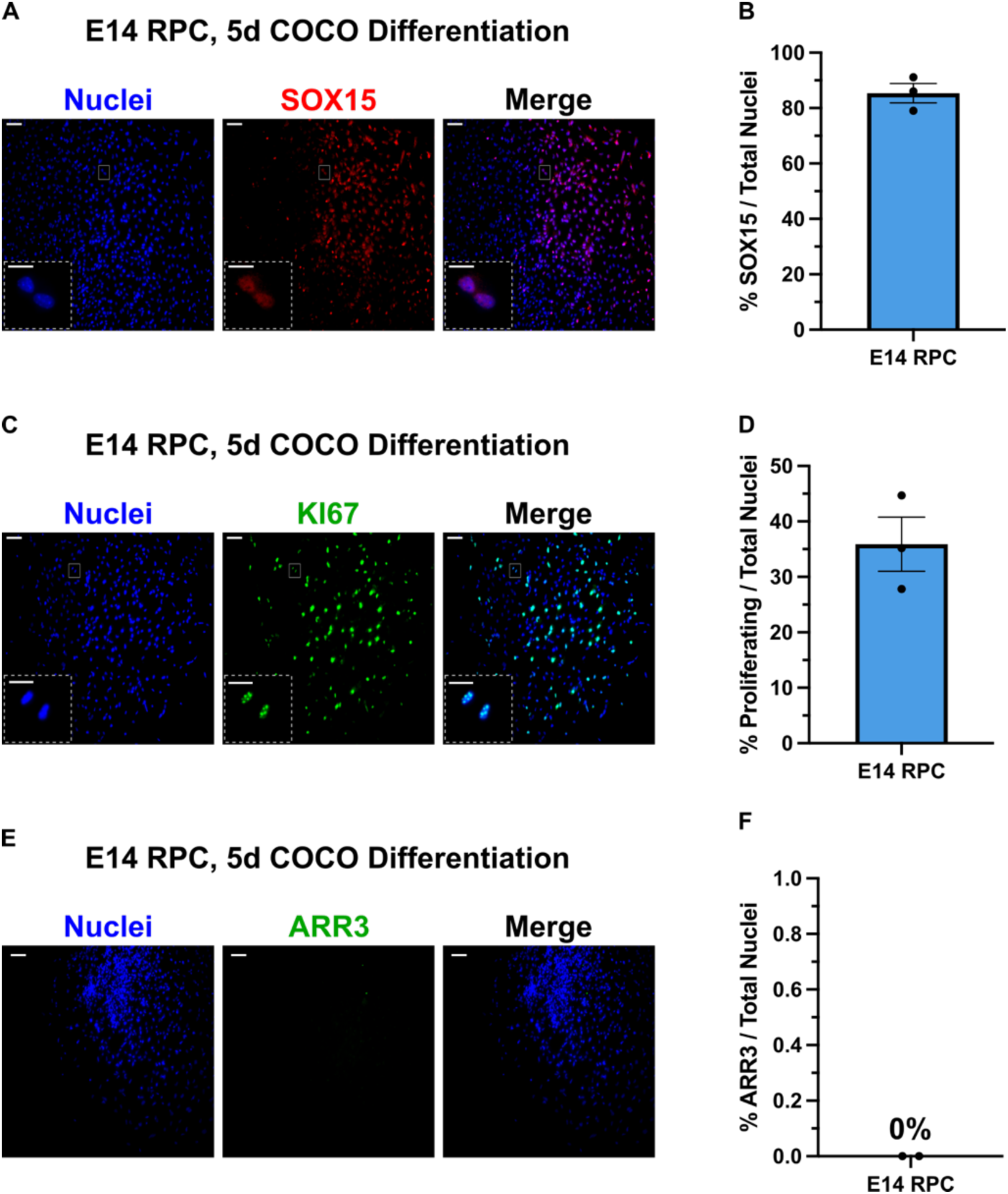
SOX15 is expressed at the protein level in proliferating RPCs after 5 days of COCO exposure. **(A)** Representative immunofluorescent micrographs of E14 RPC clones differentiated in COCO for 5 days and stained for SOX15. **(B)** Quantification of the percentage of total cells in the clone that express SOX15 (described in **A**). *N* = 3 clones. **(C)** Representative immunofluorescent micrographs of E14 RPC clones differentiated in COCO for 5 days and stained for proliferation marker KI67. **(D)** Quantification of the percentage of total cells in the clone that express KI67 (described in **A**). *N* = 3 clones. **(E)** Representative immunofluorescent micrographs of E14 RPC clones differentiated in COCO for 5 days and stained for ARR3. **(F)** Quantification of the percentage of total cells in the clone that express ARR3 (described in **E**). *N* = 2 clones. Error bars represent mean ± SEM. For all micrographs main image scale bar is 100 μm and insert scale bar is 25 μm.

### SOX15 is expressed in a subset of early embryonic RPCs *in vivo*

To ensure that *Sox15* upregulation in the presence of COCO was not an artifact of our *in vitro* tissue culture system, we first looked for *Sox15* mRNA expression in the developing and adult murine retina using RNAscope fluorescence *in situ* hybridization. At E14, *Sox15* was expressed widely throughout the retina in both the neuroblast layer (NBL, where RPCs reside) and the newly postmitotic ganglion cell layer (GCL). When compared to the broad RPC marker *Chx10*, *Sox15* mRNA puncta were both less numerous within individual cells and there were fewer overall numbers of *Sox15*-positive cells. Most important, there was a subset of *Chx10*-positive RPCs in the outer NBL that were also *Sox15*-positive, as this is a region of the developing retina where newly differentiating photoreceptors will reside (Fig. 5A). At E19, when RPCs have mostly stopped producing cone photoreceptors, there was markedly reduced numbers of *Sox15*-positive cells with much less overlap with *Chx10*-positive RPCs when compared to E14. There also was sparse *Sox15* labelling in the GCL and in the outer nuclear layer (ONL), where differentiated retinal ganglion cells and photoreceptors reside, respectively (Fig. 5B). As in the embryonic retina, in the adult there was sparse but widespread presence of the *Sox15* transcript, with the most numerous being in the GCL. Very few photoreceptors in the ONL were *Sox15*-positive and there was only one small punctum per cell, indicating low expression (Fig. 5C). Staining for SOX15 protein largely corresponded with the *in situ* hybridization data. At E12, a large proportion of CHX10-positive RPCs were also SOX15-positive (Fig. 6A) and by E14 there was a reduction in SOX15-positive cells with a smaller subset of CHX10 RPCs co-staining in the NBL. Most cells in the newly formed GCL were also positive for SOX15 (Fig. 6B). At E19 there was a substantial reduction in SOX15-positive cells in the NBL with more CHX10-negative SOX15 cells and persistent high expression in the GCL (Fig. 6C). By early post-natal and adult timepoints, SOX15 protein was restricted to a subset of retinal ganglion cells (Fig. 6D,E). Despite there being low levels of *Sox15* transcript in the adult ONL and inner nuclear layer (INL), these seemed to be translationally repressed when looking at SOX15 protein staining. In total, these *in vivo* expression data suggest that the subset of RPCs expressing SOX15 in the E14 developing retina (Fig. 5A; Fig. 6B) correspond to the *in vitro* d5 COCO treated RPCs that have a spike in *Sox15* expression in our RNA-seq experiment (Fig. 4).

**Fig. 5.**
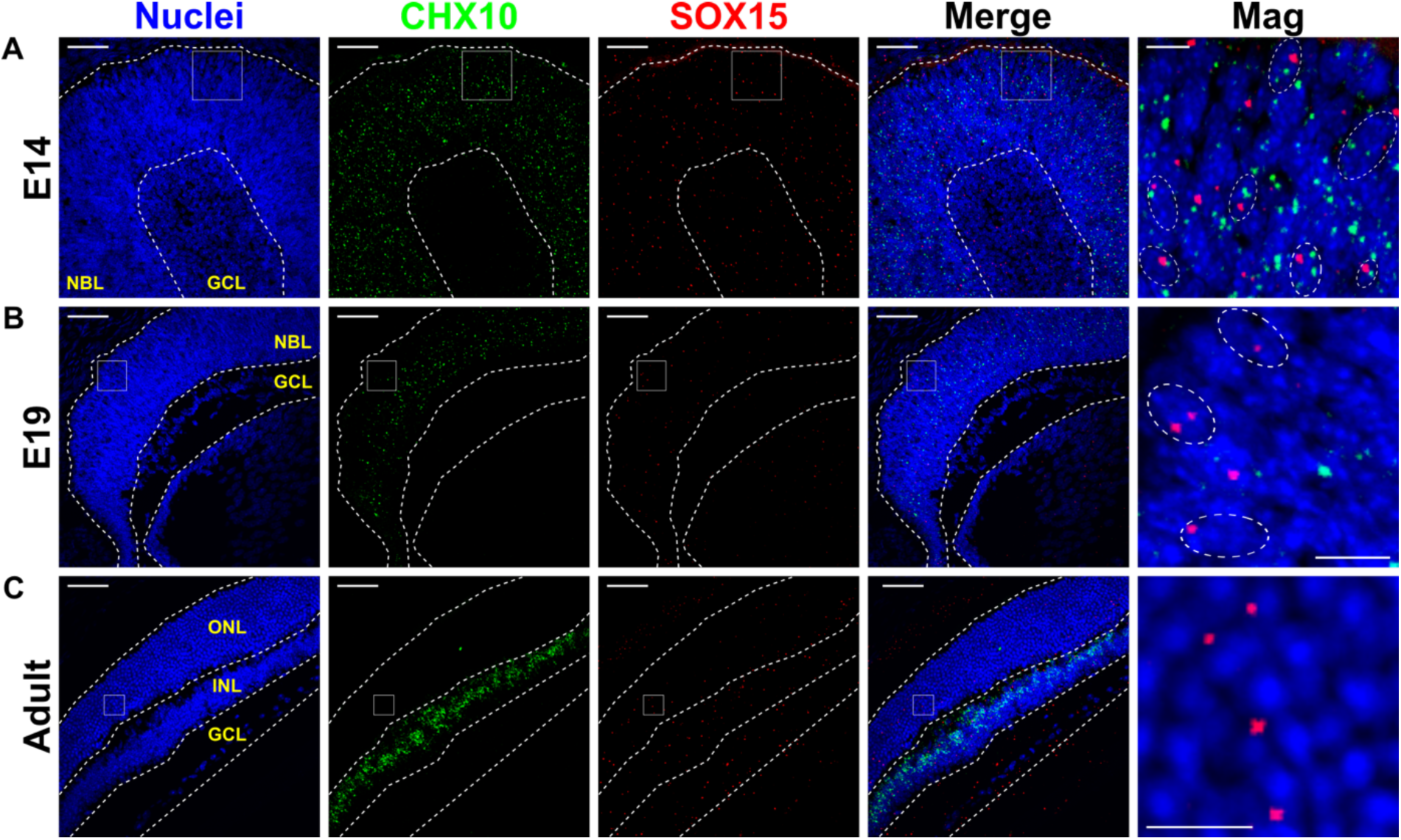
*Sox15* transcript is expressed *in vivo* in embryonic RPCs. **(A-C)** Representative immunofluorescent micrographs of RNA-scope *in situ* hybridization with probes for *Chx10* and *Sox15* on sections from embryonic and adult retinas. *Chx10* is a broad RPC marker embryonically and a bipolar cell marker postnatally. E14 dotted outlines in magnified (Mag) image show putative colocalization of *Sox15* and *Chx10* in single RPCs and E19 show *Sox15* in *Chx10*-negative putatively post-mitotic non-RPC cells. Adult magnified image shows sparse *Sox15* expression in the photoreceptor layer. NBL (neuroblast layer), GCL (ganglion cell layer), ONL (outer nuclear layer), INL (inner nuclear layer). *N* = 3 eyes per timepoint. For all micrographs main image scale bar is 50 μm and magnified scale bar is 10 μm.

**Fig. 6.**
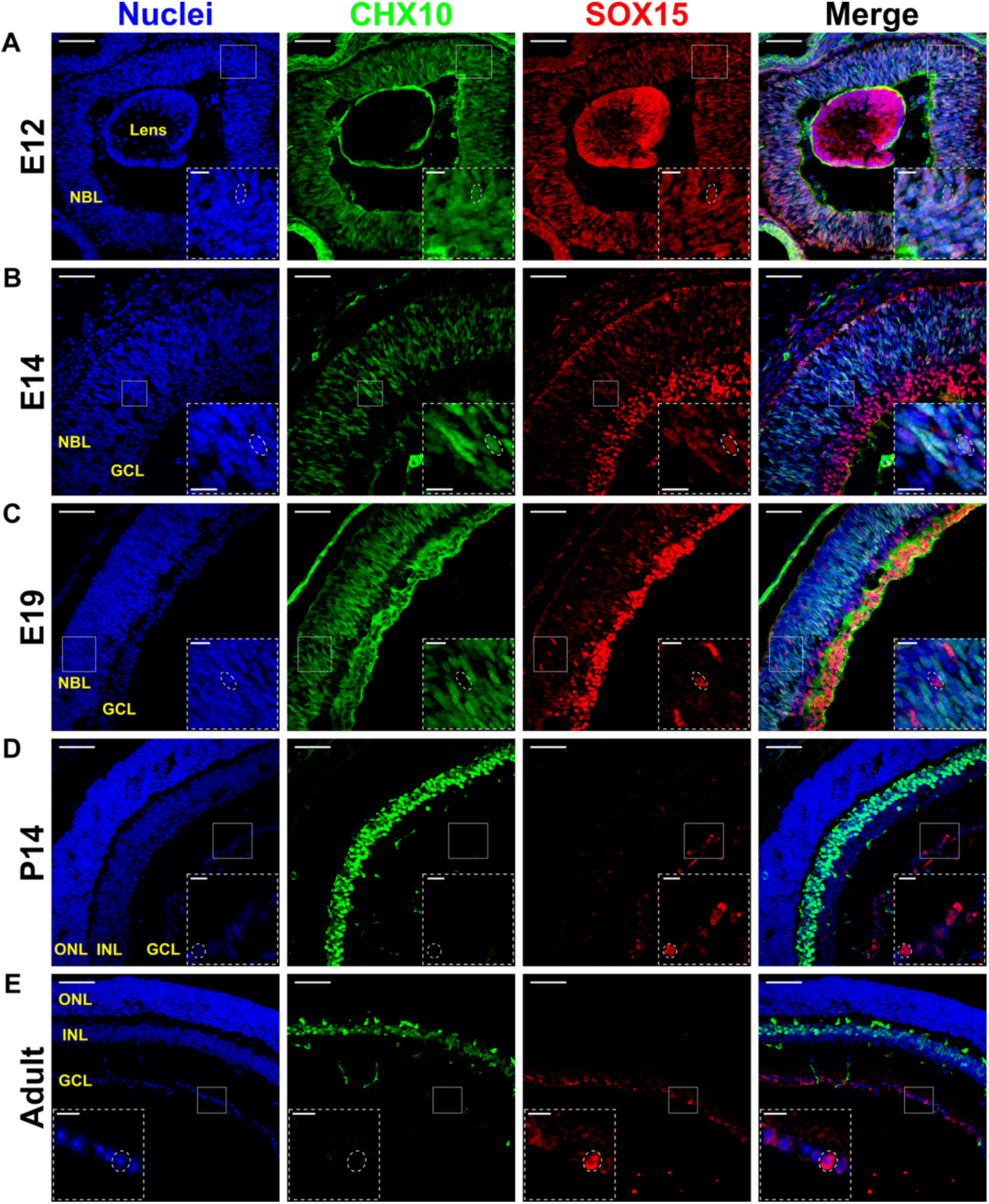
SOX15 protein is expressed *in vivo* in embryonic RPCs and early postnatal and adult retinal ganglion cells. Representative immunofluorescent micrographs of retinal sections stained with CHX10 and SOX15. **(A,B)** E12 and E14 magnified inserts show an outlined RPC co-expressing CHX10 and SOX15 in the neuroblast layer. **(C)** E19 magnified insert shows an outlined presumptive post-mitotic cell expressing SOX15 but not CHX10 in the neuroblast layer. **(D,E)** Postnatal day 14 (P14) and adult magnified inserts show an outlined retinal ganglion cell expressing SOX15. *N* = 3 eyes per timepoint. For all micrographs main image scale bar is 50 μm and magnified insert scale bar is 25 μm.

### *Sox15* knockdown causes a reduction in colony formation from embryonic RPCs

Based on the RNA-seq experiment and *in vivo* expression of SOX15, we predicted that SOX15 may be involved in promoting a cone-restricted differentiation fate in RPCs from the embryonic retina. However, we first wanted to test if there was an effect on proliferation or survival of RPCs and RSCs prior to initiation of differentiation using siRNA knockdown (KD) with a colony forming sphere assay. After 6 days of colony formation, *Sox15* KD caused a severe reduction in colony number when compared to a non-targeting (NT) siRNA or the siRNA delivery medium alone (NT: 6.250 colonies ± 0.8624 versus Sox15: 0.8889 ± 0.2271 colonies; Fig. 7A,B). This effect also was seen to a lesser extent in RPCs from the E19 retina (NT: 6.0 colonies ± 0.4075 versus Sox15: 3.167 colonies ± 0.5081; Fig. 7A,C). Most interesting, there was no effect on colony formation when the initiating cell was an RSC, compared to an RPC; and this was true of RSCs derived from both the embryonic and adult retina (Fig. 7D,E). qPCR confirmed that the siRNAs were causing an ∼90% reduction in *Sox15* expression in RPCs (Fig. 7F). There was a high degree of variability in *Sox15* expression in the NT group (Fig. 7F), however, this was not reflected in actual sphere colony numbers (Fig. 7B,C). The lack of difference in colony number between delivery medium only and NT groups suggests that elevating levels of *Sox15* expression above baseline (as seen in NT group qPCR) does not result in increased colony formation. It is conceivable that the reduction in RPC colony formation with *Sox15* KD is caused by either an inhibition of proliferation or a lack of cell survival. To test this, we measured colony formation at both day 3 and 6. From both E14 and E19 RPCs, there was no significant difference in colony formation at day 3 between NT siRNA and Sox15 siRNA, however, there was significant reduction in colony formation by day 6 (Fig. 7G,H). These data suggest that *Sox15* KD reduces cell survival rather than proliferation, as the E14 colonies that formed by day 3 would not be expected to disappear by day 6 if proliferation were inhibited (Fig. 7G). Even if the presence of E14 colonies at day 3 was due to a delay in siRNA effect, an inhibition of proliferation should not cause such a reduction in already formed colonies by day 6. It is apparent that E19 RPCs have a longer delay in initiating proliferation, as both NT and Sox15 siRNA groups showed no colony formation at day 3 (Fig. 7H). A proliferation reduction in E19 RPCs is not consistent with the data because one would not expect any colonies to be able to form by day 6 with *Sox15* KD, whereas we do see a small number of colonies compared to the much larger NT group (Fig. 7H). Given that *Sox15* has proposed regulatory interactions with WNT signaling (Moradi et al., 2017), one other possibility we did not explore is a reduction in cell adhesion causing the reduction in colony forming ability. One limitation to the colony forming assay is that only a small percentage of the total cells plated form a colony over the 6-day growth period. Therefore, the effects of *Sox15* KD we are assaying are necessarily on the most highly proliferative progenitors that can form a sphere colony and not progenitors that may only undergo a few cell divisions. Collectively, a lack of RPC cell survival with *Sox15* KD can explain both the E14 and E19 observations.

**Fig. 7.**
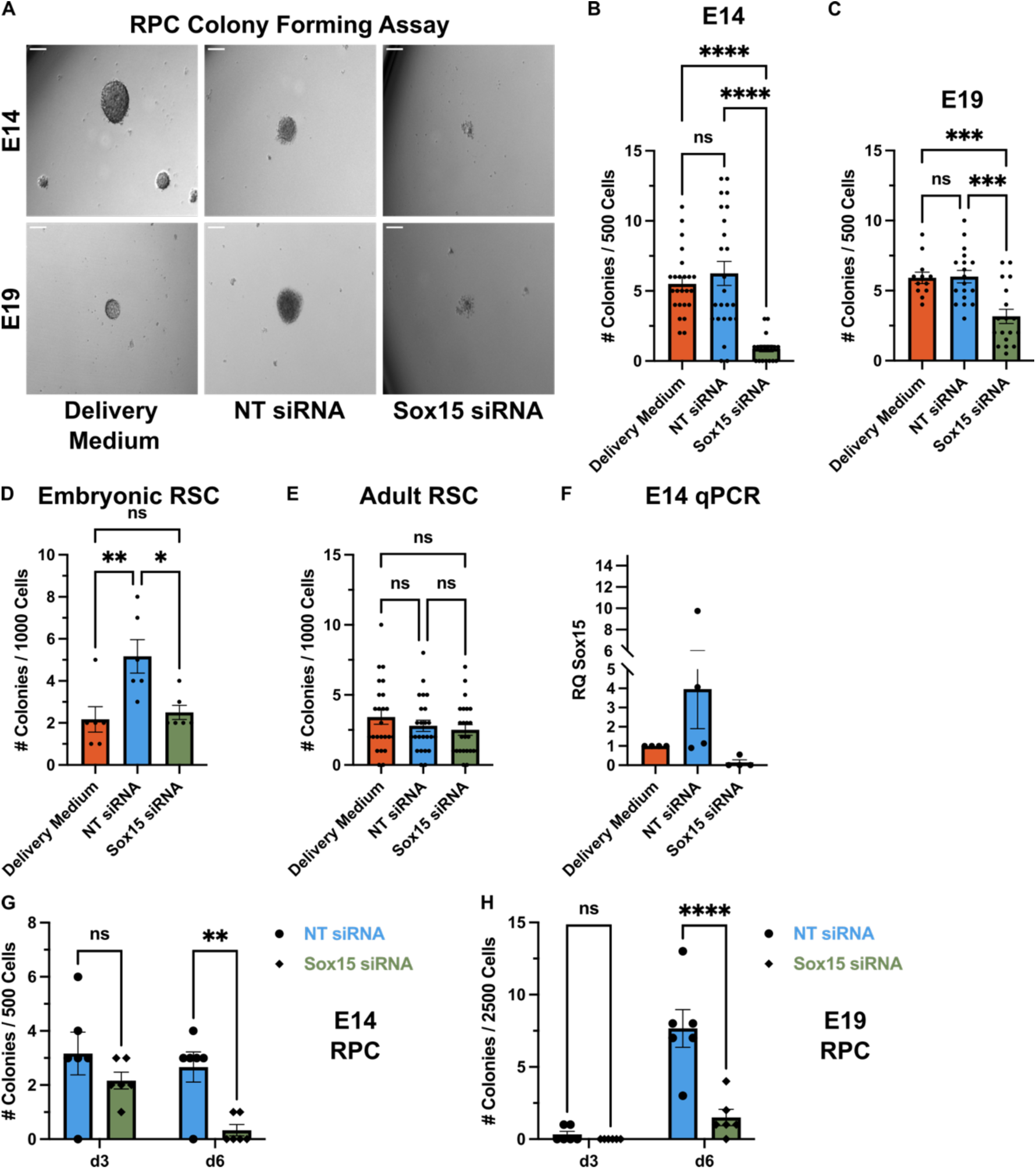
*Sox15* knockdown reduces the colony forming ability of embryonic RPCs but not RSCs. **(A)** Single cell suspensions of embryonic neural retina or ciliary epithelium (or adult ciliary epithelium) were plated free-floating at a clonal density (5 cells / μL) and supplemented with Accell siRNA reagents. The number of colonies were quantified after 6 days of growth. siRNAs were either non-targeting (NT) or a combined pool of 4 siRNAs targeting multiple regions of the *Sox15* gene. Representative brightfield micrographs show E14 and E19 RPC colonies after 6 days of growth. **(B)** Quantification of E14 RPC colony formation, as described in **A**. Delivery medium, *n* = 24 wells; NT siRNA, *n* = 24 wells; Sox15 siRNA, *n* = 18 wells; one-way ANOVA with Tukey’s multiple comparisons test, *\*\*\*\*p* <0.0001; ns, *p* = 0.6534. **(C)** Quantification of E19 RPC colony formation, as described in **A**. Delivery medium, *n* = 12 wells; NT siRNA, *n* = 18 wells; Sox15 siRNA, *n* = 18 wells; one-way ANOVA with Tukey’s multiple comparisons test, *\*\*\*p* = 0.0009 (delivery vs. Sox15), *\*\*\*p* = 0.0001 (NT vs. Sox15), ns, *p* =0.9924. **(D)** Quantification of E14 RSC colony formation, as described in **A**. Delivery medium, *n* = 6 wells; NT siRNA, *n* = 6 wells; Sox15 siRNA, *n* = 6 wells; one-way ANOVA with Tukey’s multiple comparisons test, *\*p* = 0.0187, *\*\*p* = 0.0086, ns, *p* = 0.9207. **(E)** Quantification of adult RSC colony formation, as described in **A**. Delivery medium, *n* = 24 wells; NT siRNA, *n* = 24 wells; Sox15 siRNA, *n* = 24 wells; one-way ANOVA with Tukey’s multiple comparisons test; ns, *p* >0.05. **(F)** qPCR of E14 colonies after 6 days of growth and siRNA supplementation. Fold change (RQ) was measured against the delivery medium control (calibrator) and *Actb* and *Gapdh* were used as endogenous control genes. *N*= 3 experiments. **(G, H)** Quantification of E14 and E19 RPC colony formation (as described in **A)** after 3 and 6 days of growth. *N*= 6 wells for each timepoint and condition. Two-way ANOVA with Sidak multiple comparison test; E14, *\*\*p*= 0.0094; E19, *\*\*\*\*p* > 0.0001. Error bars represent mean ± SEM. For all micrographs scale bar is 100 μm.

### *Sox15* promotes cone photoreceptor differentiation in RPCs through inhibition of rod photoreceptor differentiation

We next tested the effects of *Sox15* KD on directed photoreceptor differentiation. As with experiments above, we differentiated individual clonally derived spheres in PAN (pan retinal), COCO (cone promoting), or T + RA (rod promoting) conditions with lentiviral (LV) delivery of shRNAs. We chose to use adult RSC colonies for these functional experiments because we found that it was easier to get high infection rates in adult RSC colonies compared to embryonic RPC colonies, and others have found also that LVs do not infect RPCs in the developing retina effectively (Gurdita et al., 2023). We reasoned also that the lack of a proliferation or survival effect on RSC colonies with *Sox15* KD (compared to embryonic RPCs) would allow us to directly interrogate differentiation effects in these cultures. To ensure the effectiveness of our LV shRNA, we stained mESC transduced colonies with SOX15. Pluripotent mESCs ubiquitously and consistently express SOX15, making them an ideal system to observe a potential reduction in protein levels (Moradi et al., 2017). With LV delivery of Sox15 shRNA, there was a significant reduction in fluorescent intensity compared to a scrambled (Scr) shRNA control (Scr: 50.71 ± 4.276 versus Sox15: 35.85 ± 2.902; Fig. S3A,B). The mESC colony staining also confirms the specificity of the Sox15 antibody used in Figure 6. Differentiation condition (PAN, COCO, or T + RA) did not affect transduction efficiency, as there were similarly high percentages of infected cells at the end of the differentiation period (PAN: 79.13% ± 5.170 versus COCO: 64.72% ± 8.272 versus T + RA: 57.83% ± 9.810; Fig. S4A,B).

We predicted that a reduction in *Sox15* expression should cause a reduction in cone photoreceptor numbers in each clone, however, we observed an effect only on rod photoreceptor number. Under PAN conditions, *Sox15* KD caused a large (4.6-fold) increase in the percentage of RHO-positive cells in each clone (Scr: 7.767% ± 0.9539 versus Sox15: 35.73% ± 5.061; Fig. 8A,B), and unexpectedly, there was no change in the percentage of ARR3-positive cells with *Sox15* KD (Fig. 8C). We saw similar effects in COCO conditions, where there was a significant increase in the percentage of RHO-positive cells in each clone with *Sox15* KD and no effect on ARR3-positive cone cells (Scr RHO: 8.80% ± 0.9055 versus Sox15 RHO: 20.64% ± 4.698; Fig. 8D-F). The RHO increase in COCO conditions was 2.3-fold, which is considerably smaller than PAN conditions, and this is likely because COCO causes such a large bias in cone differentiation under control conditions (Fig. 8C versus 8F; 2.6-fold increase in ARR3 with COCO versus PAN). In T + RA there was no increase in RHO or ARR3 percentage with *Sox15* KD (Fig. 8G-I). The fact that there is no T + RA increase in rods with *Sox15* KD is likely due to a ceiling effect of rod differentiation in response to T + RA. As mentioned, RSC colonies contain a mixture of both RPC and RPE progenitors, the latter of which does not develop into neural retinal cell types like photoreceptors (Khalili et al., 2018; Baakdhah et al., 2021). Therefore, there is likely no more availability for increased rod differentiation even with *Sox15* KD, as RPE progenitors cannot differentiate into rods and T + RA has such a strong instructive effect on RPC differentiation (up to 80% of the cells in the clone can be RHO-positive; Fig. 8H). In T + RA, it is anticipated also that there would be no effect on ARR3 numbers with shRNA transduction (Fig. 8I). Of the three differentiation conditions, T + RA exhibits the lowest amount of cone differentiation, and our group has previously shown that in a combined T + RA and COCO treatment, T + RA is inhibitory to COCO activity (Khalili et al., 2018).

**Fig. 8.**
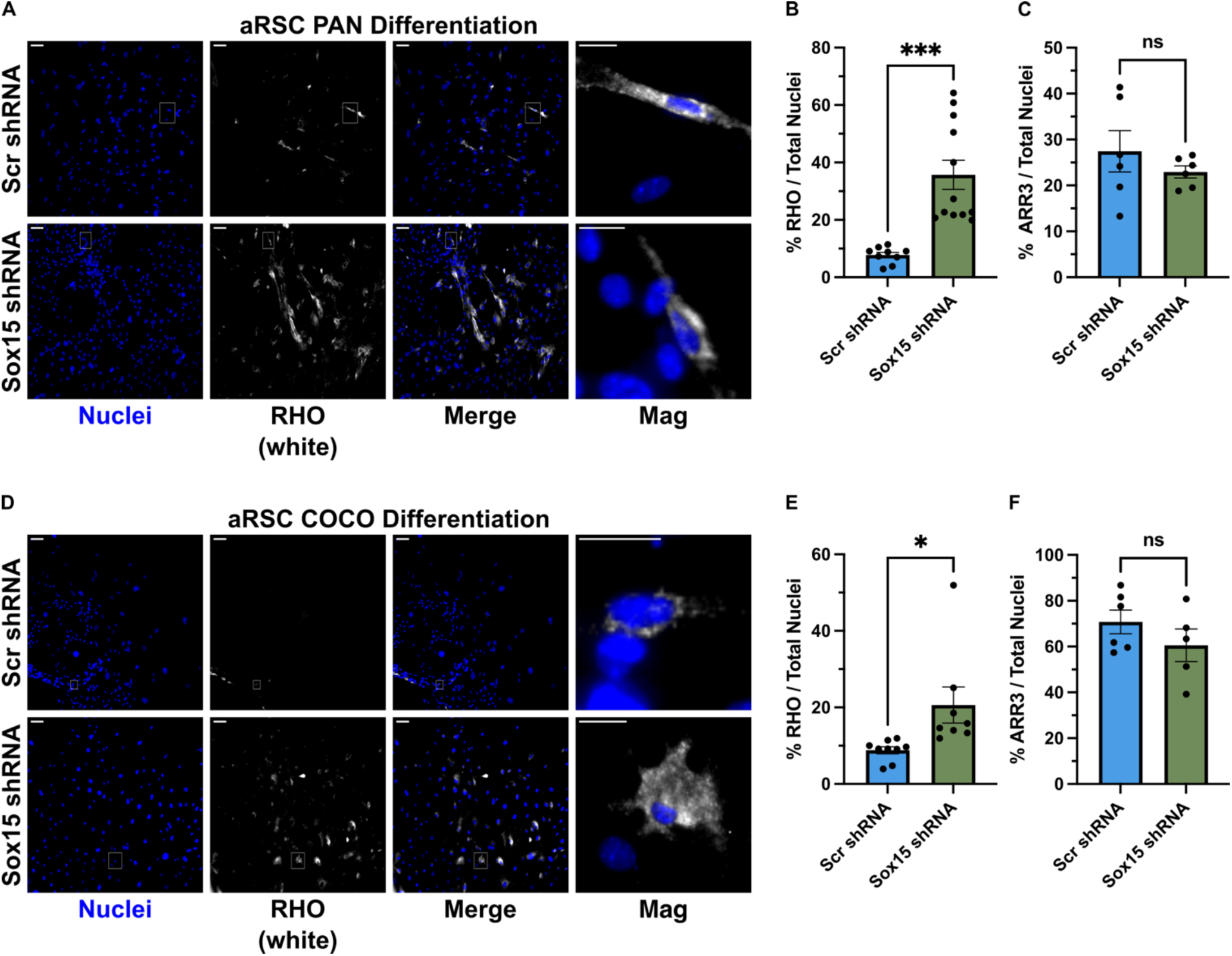
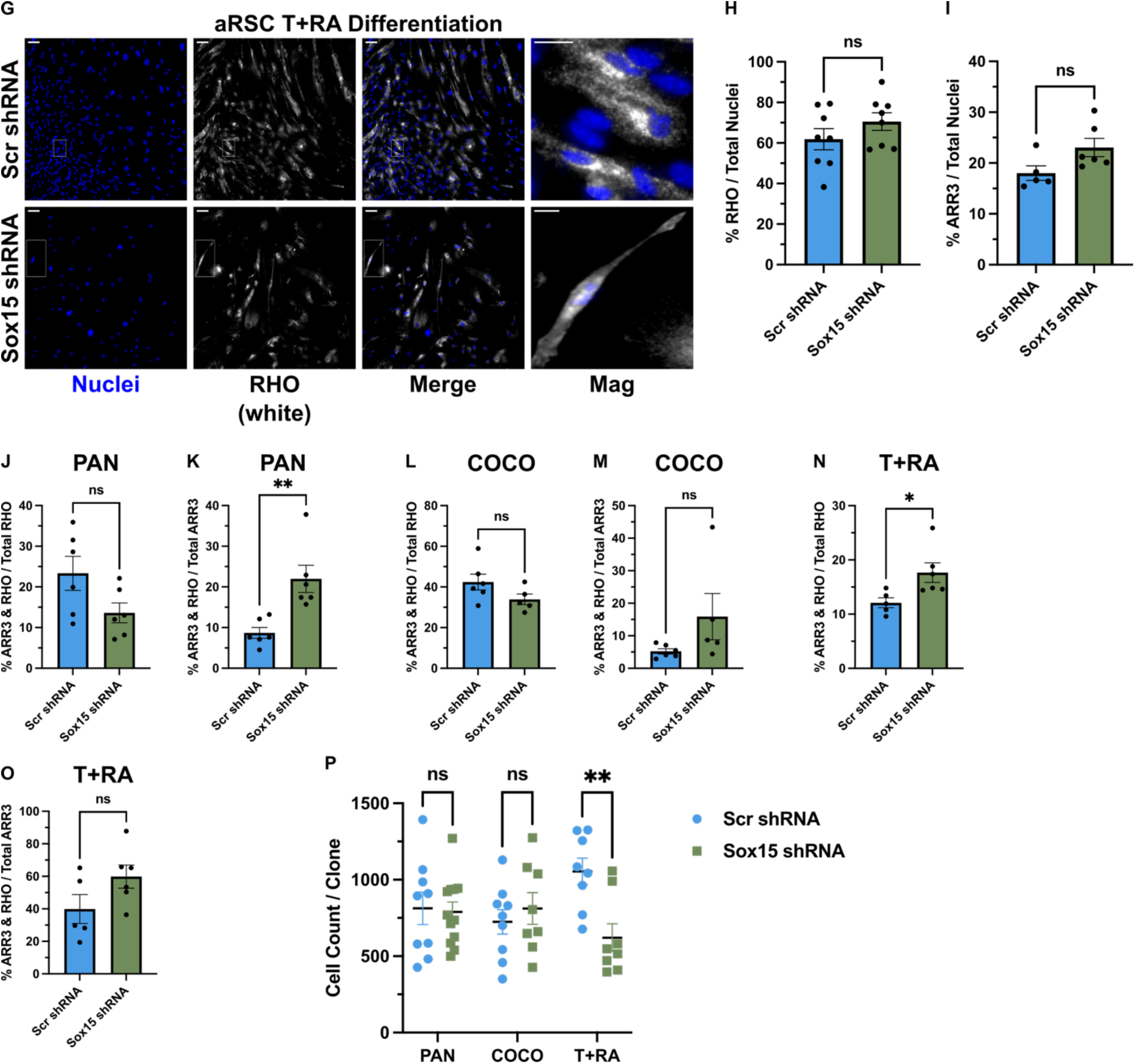
*Sox15* inhibits rod photoreceptor differentiation in RPCs. Adult RSC-derived sphere colonies were plated at a single sphere colony per well in PAN, COCO, or T + RA differentiation conditions for 2 weeks and then stained for RHO or ARR3. 2 days after plating cells were transduced with scrambled (Scr) or Sox15 shRNA LVs. **(A)** Representative immunofluorescent micrographs of RHO expressing cells in the clone after PAN differentiation and shRNA LV transduction. **(B)** Quantification of the percentage of total cells that are RHO-positive in each clone (described in **A**). Scr shRNA, *n* = 9 clones; Sox15 shRNA, *n* = 12 clones; two-tailed unpaired Student’s t-test, *\*\*\*p* = 0.0002. **(C)** Quantification of the percentage of total cells that are ARR3-positive in each clone (described in **A**). Scr shRNA, *n* = 6 clones; Sox15 shRNA, *n* = 6 clones; two-tailed unpaired Student’s t-test, ns, *p* = 0.3597. **(D)** Representative immunofluorescent micrographs of RHO expressing cells in the clone after COCO differentiation and shRNA LV transduction. **(E)** Quantification of the percentage of total cells that are RHO-positive in each clone (described in **D**). Scr shRNA, *n* = 9 clones; Sox15 shRNA, *n* = 8 clones; two-tailed unpaired Student’s t-test, *\*p* = 0.0192. **(F)** Quantification of the percentage of total cells that are ARR3-positive in each clone (described in **D**). Scr shRNA, *n* = 6 clones; Sox15 shRNA, *n* = 5 clones; two-tailed unpaired Student’s t-test, ns, *p* = 0.2660. **(G)** Representative immunofluorescent micrographs of rhodopsin expressing cells in the clone after T + RA differentiation and shRNA LV transduction. **(H)** Quantification of the percentage of total cells that are RHO-positive in each clone (described in **G**). Scr shRNA, *n* = 8 clones; Sox15 shRNA, *n* = 8 clones; two-tailed unpaired Student’s t-test, ns, *p* = 0.2185. **(I)** Quantification of the percentage of total cells that are ARR3-positive in each clone (described in **G**). Scr shRNA, *n* = 5 clones; Sox15 shRNA, *n* = 6 clones; two-tailed unpaired Student’s t-test, ns, *p* = 0.0638. **(J)** Quantification of the percentage of total RHO-positive cells in each PAN differentiated clone that are ARR3 and RHO positive. Scr shRNA, *n* = 6 clones; Sox15 shRNA, *n* = 6 clones; two-tailed unpaired Student’s t-test, ns, *p* = 0.0721. **(K)** Quantification of the percentage of total ARR3-positive cells in each PAN differentiated clone that are ARR3 and RHO positive. Scr shRNA, *n* = 6 clones; Sox15 shRNA, *n* = 6 clones; two-tailed unpaired Student’s t-test, *\*\*p* = 0.0042. **(L)** Quantification of the percentage of total RHO-positive cells in each COCO differentiated clone that are ARR3 and RHO positive. Scr shRNA, *n* = 6 clones; Sox15 shRNA, *n* = 5 clones; two-tailed unpaired Student’s t-test, ns, *p* = 0.1161. **(M)** Quantification of the percentage of total ARR3-positive cells in each COCO differentiated clone that are ARR3 and RHO positive. Scr shRNA, *n* = 6 clones; Sox15 shRNA, *n* = 5 clones; two-tailed unpaired Student’s t-test, ns, *p* = 0.1332. **(N)** Quantification of the percentage of total RHO-positive cells in each T + RA differentiated clone that are ARR3 and RHO positive. Scr shRNA, *n* = 5 clones; Sox15 shRNA, *n* = 6 clones; two-tailed unpaired Student’s t-test, *\*p* = 0.0301. **(O)** Quantification of the percentage of total ARR3-positive cells in each T + RA differentiated clone that are ARR3 and RHO positive. Scr shRNA, *n* = 5 clones; Sox15 shRNA, *n* = 6 clones; two-tailed unpaired Student’s t-test, ns, *p* = 0.1108. **(P)** Quantification of the total number of cells per clone (from 4 images taken per clone) in each differentiation and shRNA condition. PAN: Scr shRNA, *n* = 9 clones; Sox15 shRNA, *n* = 12 clones. COCO: Scr shRNA, *n* = 9 clones; Sox15 shRNA, *n* = 8 clones. T + RA: Scr shRNA, *n* = 8 clones; Sox15 shRNA, *n* = 8 clones. Two-way ANOVA with Sidak’s multiple comparisons test; *\*\*p* = 0.0053; ns (*p* >0.05). Error bars represent mean ± SEM. For all micrographs, magnified images show examples of a single RHO-positive cell, main image scale bar is 100 μm, and magnified scale bar is 25 μm.

It was surprising that the increases in rod differentiation with *Sox15* KD did not cause a concomitant decrease in cone differentiation in PAN and COCO conditions. We predict that the same RPCs are giving rise to rods and cones, and that it is primarily exposure to cell-extrinsic niche factors that cause them to become restricted to a specific photoreceptor type. There are, however, mutations in genes such as *Nr2e3* that can cause an instability in cell type, with rod-cone hybrid photoreceptor identity in both gene expression and cellular features (Corbo and Cepko, 2005). Therefore, we questioned whether *Sox15* KD may be causing an instability of photoreceptor identity through co-staining with rod and cone markers. Under PAN differentiation, there was no significant difference in the percentage of total RHO-positive cells that were co-expressing RHO and ARR3 (Fig. 8J), but there was a significant increase with *Sox15* KD in the percentage of total ARR3-positive cells that were double positive (Scr: 8.683% ± 1.347 versus Sox15: 21.97%. ± 3.339; Fig. 8K). In a similar vein, with COCO treatment there was no significant difference in percentage of total RHO-positive cells that were RHO and ARR3 double positive with *Sox15* KD (Fig. 8L). There was higher variability in COCO *Sox15* KD when looking at percentage of total ARR3 that were double positive, and so the average increase in *Sox15* KD did not reach statistical significance (Scr: 5.217% ± 0.8052 versus Sox15: 15.88% ± 7.089; Fig. 8M). However, there were some clones that had much higher percentages of double positives and the average fold increase was similar between PAN and COCO (Fig. 8K versus 8M; 2.5-fold versus 3-fold). Finally, T + RA conditions showed significant *Sox15* KD increase in the double positive cells as a percentage of total RHO (Scr: 12.10% ± 0.9209 versus Sox15: 17.65% ± 1.802; Fig. 8N) and no significant difference in double positive percentage of total ARR3 (Fig. 8O). Overall, cone-rod hybrid cells (RHO and ARR3 positive) make up a minority of total rod or cone photoreceptors in any differentiation condition, and the increases in rod differentiation in PAN and COCO conditions with *Sox15* KD cannot be accounted for wholly by increased numbers of hybrid cells. In fact, if double-positive hybrid cells are removed from the mean RHO percentages, there are slightly higher fold increases in rods with PAN and COCO (Fig. S5A,B). While it did appear that *Sox15* KD did not influence cone differentiation, when considering double-positives in PAN and COCO there are higher reductions in true cones (ARR-positive only) with *Sox15* KD (Fig. 8K,M; Fig. S5C,D). Similarly, *Sox15* KD in T + RA actually caused a greater decrease in true cones that were not double positive and no fold change in true rods (Fig. S5E,F). Taken together, these data suggest that *Sox15* promotes cone differentiation through inhibiting rod differentiation programs. When knocked down, cells that would have differentiated to cones now fully differentiate to rods, and a smaller number of cells do not fully complete rod differentiation. We hypothesize that with more differentiation time these hybrid photoreceptors (expressing a hybrid of rod and cone markers) would resolve to become fully differentiated rods.

Lastly, we examined if *Sox15* KD had any potential effect on proliferation or survival in our differentiation conditions through observing overall clone size at the end of the differentiation period. Previous data from our group has shown that directed photoreceptor differentiation through COCO or T + RA is likely due only to an effect on cell fate and not proliferation or survival (Khalili et al., 2018). There were no differences in total clone size with *Sox15* KD in PAN or COCO, but surprisingly a large reduction in clone size in T + RA (Scr T + RA: 1054.875 cells ± 86.370 versus Sox15 T + RA: 621.000 cells ± 90.834; Fig. 8P). Given the lack of evidence for a proliferation effect with *Sox15* KD shown in Figure 7, and no indication of survival effects in PAN or COCO clones, we hypothesize that this is due to a differentiation effect. We conjecture that the combination of rod differentiation promoting effects with T + RA and *Sox15* KD causes an enhanced speed of rod differentiation, and therefore less time for RPCs to initially proliferate before going post-mitotic. In the future it will be important to test multiple time points in this differentiation paradigm with *Sox15* KD.

## Discussion

In this study, we explored a fundamental question in developmental neurobiology: what are the cellular and molecular mechanisms progenitors use to generate cellular diversity in the nervous system? Key to this question are two general paradigms: one where there are predetermined and fate-restricted progenitors for different cell classes and types, and another where there are non-restricted multipotential progenitors that pass through temporal competence periods where they can generate a host of different cell types (Livesey and Cepko, 2001; Cepko, 2014; Lodato and Arlotta, 2015). The former strategy may also be referred to as lineage dependent and the latter lineage independent. These two strategies can be broken down further into what extent cell extrinsic versus intrinsic factors contribute to progenitor cell lineage (Livesey and Cepko, 2001; Cayouette et al., 2006; Cepko, 2014). We chose to focus on the developing neural retina to probe these questions, as it has been a historically fruitful developmental neurobiology model that provides many technical advantages (Bassett and Wallace, 2012; Cepko, 2014; Zhang et al., 2023). Further, we focused in on the specific retinal lineage of cone photoreceptor differentiation from RPCs. This is a particularly interesting lineage to explore for both translational reasons (Fritsche et al., 2014), and because cone densities across the retina can vary widely between species (Choi et al., 2024) despite a high degree of evolutionary conservation in the known molecular elements that underly cone photoreceptor identity (Swaroop et al., 2010; Brzezinski and Reh, 2015).

The data presented here favor a model where cone photoreceptors are produced from RPCs that are restricted in their competence, the mechanism of which is mediated by the cell-extrinsic retinal niche factor COCO. The RPCs giving rise to cones are not, however, predetermined to cone-restricted fate, as the same progenitors when differentiated in the absence of COCO do not exhibit cone-only differentiation. Moreover, the ability to become cone-restricted does not follow a strict temporal competence model. When RPCs in the current study were derived from a late embryonic period where cones are not normally being produced *in vivo* (Young, 1985; Rapaport et al., 2004), they were still able to become cone-restricted when exposed to COCO. Therefore, cone photoreceptor differentiation is not dependent on a predetermined cell intrinsic lineage, and the temporal competence observed *in vivo* is likely facilitated by cell extrinsic factors in a changing retinal niche. Indeed, our group and others have shown that certain retinal niche factors like taurine and retinoic acid are inhibitory to cone differentiation and promote rod photoreceptor differentiation from RPCs (Kelley et al., 1999; Young and Cepko, 2004; da Silva and Cepko, 2017; Khalili et al., 2018). It is possible that some aspects of cellular diversity are not instantiated at the level of the progenitor, and rather are realized once the cell has become post-mitotic (Lodato and Arlotta, 2015). In cone photoreceptor development, there is such a model whereby all photoreceptor subtypes start from a common post-mitotic photoreceptor precursor, and then photoreceptor subtype is subsequently determined (Swaroop et al., 2010). In contrast, our data favors the interpretation that photoreceptor specification is determined at the level of the progenitor, and this is in line with other recent reports of observable cone bias in RPCs (Cepko, 2014). We have previously shown that application of COCO at later time points, when cells are post-mitotic, does not result in cone-restricted differentiation, supporting fate decision at the level of the proliferating RPC (Khalili et al., 2018). Our RNA-seq experiments show also that there are overall transcriptomic differences early in rod or cone directed differentiation (T + RA versus COCO) when retinal cells are still in an RPC state. Taken together, we conclude that the default differentiation pathway of RPCs is to an s-cone photoreceptor. In the absence of inhibitory cell extrinsic factors (like T + RA, TGFβ, WNT, and BMP), RPCs default towards differentiating into s-cone photoreceptors. This differentiation mechanism resembles that of neural induction, where inhibition of these same signaling pathways causes pluripotent embryonic stem cells to differentiate into neural cells (Tropepe et al., 2001; Muñoz-Sanjuán and Brivanlou, 2002). From an evolutionary perspective, there is also evidence that the s-cone photoreceptor is the ancestral photoreceptor type (Kim et al., 2016). Going forward, it will be crucial to experimentally manipulate COCO levels *in vivo*, where we predict that increasing COCO expression at later embryonic timepoints should lengthen the temporal window of cone differentiation.

We hypothesized that downstream of COCO, there should be a unique molecular marker of cone-restricted RPCs. Through RNA-seq, *Sox15* was a top differentially upregulated transcription factor in RPCs that had been exposed to COCO. We followed this lead due to *Sox15* being previously underexplored in retinal biology, to the involvement of *Sox15* in anterior CNS development, and to regulatory activity of *Sox15* with signaling pathways manipulated by COCO (Mizuseki et al., 1998; Moradi et al., 2017; Choi et al., 2023). The *Sox15* transcript and protein were expressed in a subset of RPCs at embryonic time points when cones are normally being produced from RPCs *in vivo*, but at later postnatal and adult stages the protein was restricted to retinal ganglion cells (RGCs). While we did not explore its role in RGC differentiation or function, it will be interesting in the future to study whether it also has functional roles in RGC biology given that cones and RGCs both become post-mitotic early in retinal development (Young, 1985; Rapaport et al., 2004). Using siRNA, we found that *Sox15* KD caused reduced RPC survival in a colony forming assay. In addition, in photoreceptor differentiation conditions, LV delivery of Sox15 shRNA caused an increase in rod photoreceptor differentiation at the expense of some portion of cone differentiation and without an apparent effect on cell survival. While these two results may seem contradictory, we propose that the reduction in cell survival in the colony forming assay may be due to a premature rod differentiation in the RPC colonies. RPC colonies are in suspension and normally require adherence to a laminin coated surface for prolonged differentiation and survival of their photoreceptor progeny (Ballios et al., 2012; Khalili et al., 2018). These data support an interpretation where *Sox15* promotes cone differentiation from RPCs through inhibition of rod differentiation, downstream of the molecular effects of COCO. Lending support to this analysis, a cancer study predicted that *Sox15* genetically interacts with genes important for rod differentiation (Garancher et al., 2018). It is possible that rod marker increases with *Sox15* KD is due to these markers to be turned on more quickly, rather than changing the fate of RPCs. However, previous work from our lab has shown that in PAN conditions the ∼10% of rods do not increase with prolonged culture time of up to 40 days (Ballios et al., 2012). In this work, PAN conditions are where we see the largest rod increases with *Sox15* KD, and thus it is unlikely that the NT control levels will reach *Sox15* KD levels with longer culture.

In the future, we will test whether *Sox15* ablation *in vivo* has similar rod promoting effects at the expense of cones and *in vivo* lineage trace from *Sox15*-positive RPCs that are predicted to give rise to cone-enriched clones. Our experiments knocked down *Sox15* throughout the entire course of cone differentiation, but given the expression data from RNA-seq, we predict that knockdown is only necessary in RPCs prior to terminal division to elicit rod promoting effects. A recent report suggests that temporal identity factors promote cone fate in RPCs by repressing rod fate (Javed et al., 2023), and it will be interesting to determine whether *Sox15* has any regulatory interactions with these temporal identity factors. Other transcriptions factors, such as *Otx2*, *Olig2*, *Pou2f1*, *Pou2f2*, and *Blimp1* have been shown to bias cone differentiation in mitotic or early post-mitotic retinal cells (Brzezinski et al., 2010; Hafler et al., 2012; Emerson et al., 2013; Javed et al., 2020), and it will be necessary to functionally test whether *Sox15* regulates or is regulated by these factors. Finally, there is evidence that *Sox15* has compensatory roles with other *Sox* genes such as *Sox2* (Maruyama et al., 2005; Choi et al., 2023). It is possible that double KD of these genes will confer a more significant photoreceptor fate phenotype, as *Sox2* also is expressed in RPCs (Taranova et al., 2006). The data described here propose that in response to COCO, *Sox15* is upregulated in RPCs to promote cone photoreceptor fate through inhibiting rod photoreceptor fate. However, in the absence of COCO (PAN conditions), these same RPCs will differentiate into other non-photoreceptor retinal cell types (Tropepe et al., 2000; Ballios et al., 2012; Khalili et al., 2018). Therefore, there are likely other transcription factors downstream of COCO that are upregulated or downregulated to inhibit non-photoreceptor retinal differentiation programs. We postulate that *Sox15* may only become important once an RPC has committed to a photoreceptor differentiation program.

In summary, we present the hypothesis that COCO inhibition causing RPCs to become s-cone lineage-restricted is a unifying mechanism that modulates both cone distribution and density in vertebrate retinas. This becomes especially important in species, like humans, where there are cone dense regions responsible for high acuity color vision. Indeed, our data shows that COCO directed cone differentiation is a conserved mechanism in human embryonic and adult retinal stem cell *in vitro* developmental systems. We predict that in these cone dense regions, a greater concentration of COCO causes most progenitors to produce cones at the expense of other retinal cell types. Supporting this, there is recent evidence for a restricted progenitor mechanism in cone dense retinal regions and a necessity for degrading retinal niche factors that are inhibitory to cone differentiation (da Silva and Cepko, 2017; Choi et al., 2024). It is intriguing also that new research shows COCO can inhibit angiogenesis in the retina (Popovic et al., 2021). As the cone dense macular regions of the primate retina are reduced in vasculature (Bringmann et al., 2018), and COCO is present in the early embryonic neuroblast layer of the retina (Zhou et al., 2015), this lends further support to the idea that high levels of COCO are present to promote cone-restricted differentiation and inhibit blood vessel formation.

## Materials and Methods

### Mice

All use of animals was approved by the University of Toronto Animal Care Committee in accordance with the Canadian Council on Animal Care. Animals were housed in the Division of Comparative Medicine facility at the University of Toronto and kept on a 12-hour light/dark cycle with ad libitum feeding. For studies using mouse embryos, mice were bred at 6-12 weeks of age and plugs were checked every morning to determine the approximate time of conception. The following mouse strains were purchased from Jackson Laboratories: C57BL/6J (000664). Both sexes of mice were used for all experiments. For all experiments, adult mice were first anesthetized with isoflurane and then euthanized via cervical dislocation.

### Primary mouse cell isolation

All dissection were performed with tissue in cold artificial cerebrospinal fluid (aCSF: 124 mM NaCl, 5 mM KCl, 1.3 mM MgCl_2_, 2 mM CaCl_2_, 26 mM NaHCO_3_, and 10 mM D-glucose). For embryonic retinal dissection, using curved non-serrated forceps, embryos were decapitated and eyes were removed. With straight forceps, the optic nerve was removed, and retina was accessed by tearing the sclera and then separating it from the lens. Using a fire polished glass pipette, embryonic retinas were dissociated ∼60 times and passed through a 40 μm cell strainer to ensure a single cell suspension. For adult RSC dissections, as described previously (Tropepe et al., 2000; Coles et al., 2004), the outer surface of the eye was first sprayed with 70% EtOH and then enucleated into cold aCSF using curved microdissection scissors. With these scissors, the eye was then cut in half (starting from the optic nerve opening), and retina and lens were removed using curved non-serrated forceps. The remaining eye tissue was pinned down with straight forceps and a scalpel was used to cut a strip of tissue containing the ciliary body on one side and sclera on the other. The ciliary body tissue strip was then transferred to a 35-mm petri dish containing 2mL of Dispase (Corning, 354235) for 10 minutes at 37°C. The tissue is then transferred to another dish containing a modified aCSF (high Mg^2+^ (3.2 mM MgCl_2_) and low Ca^2+^ (108μM CaCl_2_)) enzyme solution with 1.33 mg/mL trypsin (Sigma, T4799), 0.67 mg/ml hyaluronidase (Sigma, H3506), and 0.2 mg/ml kynurenic acid (Sigma, K3375) for 10 minutes at 37°C. After enzymes, the ciliary epithelium is scraped away using curved non-serrated forceps and dissociated ∼30 times using a fire polished glass pipette. This dissociated cell suspension was then spun down at 0.4 RCF for 5 minutes, pellet resuspended in serum free medium (SFM) containing 1 mg/mL of trypsin inhibitor (Worthington, LS003085), and further dissociated ∼30 times using a small borehole glass pipette. This cell suspension was then spun down at 0.4 RCF for 5 minutes and resuspended in growth factor supplemented SFM for cell counting and downstream culture. A more detailed description and video guide for RSC dissection can be found on JoVE (Coles and van der Kooy, 2010).

### Primary human cell isolation

Consent was given for human eyes to be used for research and eyes were obtained from the Eye Bank of Canada (Toronto, ON) within 24 hours post-mortem. There were no age or sex restrictions for this study and all procedures using this human tissue was approved by the University of Toronto Research Ethics Board. Eyes were placed in a petri dish with aCSF and cut into halves. Tissue strips comprising the sclera on one side and ciliary body on the other were prepared as described above. The ciliary body and Bruch’s membrane were then peeled away from the sclera and Bruch’s membrane was removed from the ciliary body with microdissection scissors. Strips of ciliary body were treated to the enzymatic regime described above, but with 20-minute incubation instead of 10. Scraping and dissociation also were performed as described in the previous section.

### Primary stem and progenitor cell culture and differentiation

Unless indicated, cells were culture in a normoxic 5% CO2 incubator at 37°C. For colony formation, primary adult RSCs or embryonic RPCs were plated in suspension culture at a clonal density of 5 cells/μL for 7 days or 6 days, respectively. At this density, single RSCs or RPCs will proliferate to give rise to a free-floating spherical colony of RSCs and downstream RPCs or exclusively RPCs, respectively. Plates should not be moved for the entirety of colony formation to ensure that clonality is maintained (Coles-Takabe et al., 2008). Cells were plated in a serum free medium (SFM) containing a 1:1 mixture of DMEM (Thermo, 12100046) and F12 (Thermo, 21700075) at a final 1x concentration, 0.6% D-glucose (Sigma, G6152), 5 mM HEPES (Thermo, 15630080), 3 mM NaHCO_3_ (Thermo, 25080094), 2 mM L-Glutamine (Thermo, 25030081), 25 μg/mL insulin (Sigma, I5500), 100 μg/mL apo-transferrin (Sigma, T2252), 20 nM progesterone (Sigma, P6149), 60 μM putrescine (Sigma, P5780), 30 nM sodium selenite (Sigma, S9133), and 1% penicillin-streptomycin (Thermo, 15140122). For colony formation the SFM is supplemented with 20 ng/mL bFGF (Sigma, F0291) and 2 ng/mL heparin (Sigma, H3194). SFM should be made fresh every 3 days. For differentiation experiments cell culture treated 24 well plates (Thermo, 142475) were first coated with laminin (Millipore, L2020). Laminin was dissolved in SFM (5 μL/mL) and the solution was added to the wells for 4 hours at 37°C. Wells were then washed with PBS 3 times for 5 minutes each wash. Pan-retinal (PAN) differentiation medium contained SFM + 20ng/μL bFGF + 2 ng/μL heparin + 1% heat inactivated FBS (Gibco, 10082147), COCO differentiation medium was PAN medium + 50 ng/μL COCO (R&D, 3356-CC-025), and taurine and retinoic acid (T + RA) differentiation medium was PAN + 100 μM taurine (Sigma, T8691) and 500 nM retinoic acid (Sigma, R2625). Individual sphere colonies were picked and put in a laminin coated well (1 sphere / well) for adherent retinal differentiation in PAN, T + RA, or COCO conditions. Fresh differentiation medium was exchanged every 3 days and medium should be supplemented with fresh COCO or T + RA every exchange. Depending on the experiment, cells were differentiated for 2-6 weeks.

### Mouse embryonic stem cell (mESC) culture and retinal organoid differentiation

Unless indicated, cells were culture in a normoxic 5% CO2 incubator at 37°C. 6 well cell-culture treated plates were coated with 0.1% gelatin (Stem Cell Technologies, 07903) for 30 minutes at room temperature. mESCs were maintained in GMEM (Thermo, 11710035), 1% ESC-qualified FBS (Gibco, 16141079), 10% KSR (Gibco, 10828028), 0.1 mM (1%) NEAA (Sigma, M7145), 1 mM (1%) sodium pyruvate (Gibco, S8636), 0.1 mM 2-Mercaptoethanol (Millipore, M6250), 1% penicillin-streptomycin, and 2000 U/mL LIF (Millipore, ESG1107). Fresh medium was exchanged every 1-2 days and medium was supplemented with LIF every medium exchange. Cells were passaged once reaching 70% confluency. For passaging, cells were washed once with PBS and then incubated at 37°C for 5 minutes in TrypLE (Thermo, 12605028). After incubation, a serological pipette was used to dissociate to a single cell suspension which was then transferred to a Falcon tube topped up with at least equal volume of PBS to dilute out the TrypLE. Cells were then spun down at 0.4 RCF for 5 minutes and split 1:6 in gelatin coated plates with fresh maintenance medium. For retinal organoid culture, we followed procedures detailed in the HIPRO protocol (Chen et al., 2020). Briefly, mESCs were dissociated into a single cell suspension using TrypLE and plated at 3000 cells / well in ultra-low attachment U bottom 96 well plates (S-bio, MS-9096UZ) in retinal differentiation medium (RDM). For the first 7 days, aggregates of mESCs were cultured in a 5% CO2, 5% O2, and 37°C incubator. RDM during this period contained GMEM, 1% NEAA, 1% sodium pyruvate, and 1% KSR. Before plating mESCs, cold retinal differentiation medium was supplemented with growth factor reduced Matrigel (Corning, 354230), so that each well contained ∼20 μL of Matrigel. After 7 days, organoids were transferred to a low attachment 6 well plate (Corning, CLS3471) with ∼ 6 organoids / well. From day 7-10, organoids were culture in retinal maturation medium (RMM), which contained DMEM/F12 with GlutaMAX (Thermo, 10565042), 1% N-2 (Thermo, 17502048), 55 μM 2-ME (Thermo, 21985023) and 1% penicillin-streptomycin. For COCO conditions, 50 ng/mL of COCO was added at day 7. From day 10 onwards, organoids were cultured in RMM supplemented with 1 mM taurine, 0.5 μM 9-*cis* retinal (Millipore, R5754), and 10 ng/mL IGF1 (Thermo, PHG0071). COCO conditions contained non-supplemented RMM with 50 ng/mL COCO. Fresh medium was exchanged every other day. After 14 days, organoids were removed for qPCR.

### Human embryonic stem cell (hESC) culture and retinal differentiation

Unless indicated, cells were culture in a normoxic 5% CO2 incubator at 37°C. 6 well tissue culture dishes were coated with hESC-qualified Geltrex (Thermo, A1569601) for 1 hour at 37°C. H9 hESCs were cultured in mTeSR1 (Stemcell Technologies, 85850) maintenance medium and passaged at 70% confluency or when differentiation started to appear in individual hESC colonies. Passaging was done using the ReLeSR (Stemcell Technologies, 100-0484) protocol at a split ratio of 1:6, and cells were passaged as 50-200 μm clumps. For RSC differentiation, hESCs were first grown in maintenance medium to over-confluency (∼10-14 days) and then switched to a retinal differentiation medium containing knock-out DMEM (Thermo, 10829018), 13% KSR, 0.1% NEAA, 1% 2-ME, and 1% GlutaMAX (Thermo, 35050061). For retinal differentiation it is important that this medium does not contain bFGF and the medium was exchanged every other day. After ∼1-month, pigmented patches should start to appear, however, there is some variability to this process and it may take slightly longer for patches to appear. To obtain RSCs, pigmented patches are cut out using a pipette tip and dissociated and cultured in a colony forming sphere assay as described in “Primary human cell isolation” and “Primary stem and progenitor cell culture and differentiation”. Individual clonally derived RSC sphere colonies were differentiated as described in “Primary stem and progenitor cell culture and differentiation”.

### siRNA colony formation

Detailed protocol on the preparation of siRNA can be found on the Horizon Discovery website. A single-cell suspension of RPCs or RSCs were prepared in a serum free solution of Accell siRNA Delivery Medium (Horizon Discovery, B-005000-500) supplemented with 20 ng/mL bFGF and 2 ng/mL heparin at a final concentration of 5 cells/μL. This solution was also supplemented with 1 μM of either Accell non-targeting control pool siRNAs (Horizon Discovery, D-001910-10-20) or Sox15 SMARTPool siRNAs (Horizon Discovery, E-042792-00-0020) containing a mixture of 4 siRNA targeting *Sox15*. 100 μL of this cell solution (at 5 cells/μL) was plated in each well of a 96 well plate. After 3 days of culture, each well was supplemented with 50 μL of growth factor supplemented delivery media containing 3 μM siRNA so that that final siRNA concentration in each well was 1 μM. After 6 days of growth the number of free-floating clonal sphere colonies was quantified.

### Lentiviral preparation

The following third generation lentiviral (LV) packaging plasmids were used: pMD2.G (Addgene, #12259), pRSV-Rev (Addgene, #12253), and pMDLg/pRRE (Addgene, #12251). Scrambled shRNA (#1864) and EGFP (#19319) plasmids also were acquired from Addgene (1864). Sox15 shRNA LV plasmid was purchased from Santa Cruz Biotechnology (sc-38428-SH) and contained a mixture of 3-5 plasmids each encoding target-specific 19-25 nt (plus hairpin) shRNAs. HEK293T cells were cultured in 0.1% gelatin coated 10 cm dishes and maintained in a feeder medium containing DMEM (Thermo, 11995065), 10% FBS (Gibco, ESC qualified), and 0.5X penicillin-streptomycin (Thermo, 15070063). HEK293T cells were passaged at 70-90% confluency using a 5-minute, 37°C TrypLE incubation. Cells were passaged at least once after thawing before transfection. Fresh medium was exchanged 2-4 hours prior to transfection and HEK293Ts were transfected at 70-80% confluency in the evening using the lipofectamine 2000 reagents and protocol (Thermo, 11668019). Each 10 cm dish received the following concentrations of plasmids: 2.5 μg pMD2.G, 2.5 μg pRSV-Rev, 5ug pMDLg/pRRE, and 10 μg shRNA plasmid. The following morning (∼16-18 hours) media was exchanged and then 24 hours after that media containing virus was collected. Viral supernatant was spun down (5 minutes, 0.4 RCF) to remove cellular debris and then run through a 0.45 μm PVDF syringe filter (Sigma, SLHV033RS). Aliquots of filtered viral supernatant were stored at −80°C.

### Lentiviral shRNA knockdown

Mouse RSCs were plated at a single cell per well in 24 well plates and differentiated as described in “Primary stem and progenitor cell culture and differentiation”. After 2 days in culture (when RSC colonies have adhered to the bottom of the well), 5 MOI each of scrambled or Sox15 shRNA LV and EGFP LV (10 MOI total) was added to each well with 8 μg/mL polybrene (Sigma, TR-1003). 24 hours later fresh differentiation medium was exchanged, and the differentiation protocol progressed as described for 2 weeks. To test the effectiveness of shRNA knockdown, the abovementioned viral combinations and amounts were added to mESC cultures in maintenance conditions as described in “Mouse embryonic stem cell (mESC) culture”. mESCs were grown for 5 days after viral transduction and then fixed and stained with Sox15 antibody for imaging. To quantify SOX15 fluorescence intensity, individual mESC colonies in scrambled or Sox15 shRNA conditions were imaged using a confocal microscope with identical acquisition settings between conditions. In ImageJ, individual colonies were manually outlined, and mean fluorescence intensity (MFI) was calculated for the maximum intensity projection of each colony.

### Immunohistochemistry, immunocytochemistry, and fluorescence *in situ* hybridization

Adult or embryonic mouse eyes were immersed in cold 4% PFA (dissolved in 1x PBS) for 4 hours at 4°C on an orbital shaker. Eyes were then rinsed in 1x PBS and transferred to a 30% sucrose solution (in 1x PBS) overnight at 4°C on an orbital shaker (or until the eyes have fully sunk to the bottom of the dish). For tissue sectioning, eyes were imbedded in Tissue-Tek O.C.T Compound (Sakura, 4583) and frozen in −80°C. A cryostat set at −20°C was used to cut 10 μm sagittal sections of the eye adhered to Fisherbrand Superfrost Plus slides (12-550-15), which were then stored at −80°C. For immunohistochemistry, slides were first dried at room temperature (RT) for 30 minutes and then a hydrophobic barrier was drawn around tissue sections. Unless indicated, all proceeding steps occur at RT. Slides were rehydrated with 1x PBS for 20 minutes and then permeabilized for 20 minutes with 0.3% triton (Thermo, A16046-AE). Sections were washed 3 time in 1x PBS with one 5-minute and two 10-minute incubation intervals, and unless indicated, this is what is referred to as “washed”. After that, slides were blocked in 10 % NGS (Jackson, AB_2336990) for 1 hour and primary antibodies were diluted in 1 % NGS for overnight incubation at 4°C. The next day, slides were washed and then incubated in Alexa Fluor secondary antibodies diluted 1:400 in 1% NGS for 1 hour at RT. Slides were again washed and then incubated with 10 μg/mL Hoechst 33342 (BD Biosciences, 561908) in PBS for 10 minutes. A final wash was done, and slides were cover slipped with antifade mounting medium (Abcam, ab104135), sealed, and stored at 4°C until imaging. Sections were imaged no later than 1 month after staining. For cells in culture, they were fixed with RT 4% PFA for 15 minutes and then washed. Fixed cells were stored at 4°C and processed for immunostaining no later than 1 month after fixation. Cells were permeabilized with 0.3% triton for 10 minutes and washed. All steps from blocking onwards were carried out as described above for tissue sections. For mouse experiments, the following primary antibodies and concentrations were used: 1:2000 rabbit cone arrestin (Millipore, AB15282), 1:250 mouse rhodopsin (Millipore, MAB5316), 1:500 rabbit S-opsin (Millipore, AB5407), 1:1000 chicken GFP (Aveslabs, GFP-1010), 1:100 rabbit KI67 (Abcam, ab15580), 1:500 rabbit Sox15 (Proteintech, 25415-1-AP), and 1:100 mouse Chx10 (Santa Cruz, sc-356619). For human cell experiments, all primary antibodies listed above were cross reactive with human tissue except for an additional human specific cone arrestin antibody: rabbit 1:500 (LS Bio, LS-C368677-100). All primary antibodies were tested on retinal tissue to ensure specificity of labelling. The following Alexa Fluor antibodies were used in various combinations at a dilution of 1:400: goat anti-chicken 488 (Thermo, A-11039), goat anti-mouse 488 (Thermo, A-11001), goat anti-rabbit 488 (Thermo, A-11008) goat anti-mouse 568 (Thermo, A-11004), goat anti-rabbit 568 (Thermo, A-11011), goat anti-mouse 647 (Thermo, A3728), and goat anti-rabbit 647 (Thermo, A32733). Each experiment contained a secondary antibody negative control to account for non-specific background signal. Fluorescence *in situ* hybridization was carried out on tissues sections (fixed as described above) using RNAscope multiplex fluorescent V2 assay (ACD, 323100). Detailed protocol steps can be found in the user manual under the “Fresh-frozen sample” section. Briefly, slides were dehydrated using a series of 50%, 70%, and 100% EtOH incubations. Next, slides were incubated with H2O2 for 10 minutes at RT and then washed. Protease IV was then added for 15 minutes at RT followed by washing. The following probes were used: mouse Sox15 (ACD, 501421) and mouse Chx10 (ACD, 438341-C3). In addition, both 3-plex positive (ACD, PN 320881) and negative (ACD, PN 320871) control probes were used to ensure the specificity of the system. TSA plus fluoresceine (Akoya, NEL741001KT) and Cy3 (Akoya, NEL744001KT) were used to fluorescently visualize probes. Details on probe incubation steps can be found in chapter 4 of the manual. Tissue sections were stored in the dark at 4°C and imaged no later than 2 weeks after hybridization.

### Microscopy and image quantification

*In vitro* cells were imaged using a Zeiss AxioObserver D1 inverted microscope with Zeiss AxioVision software. For retinal differentiations, single sphere colonies generally spread out in an adherent circular colony. This differentiated colony was split into approximately 4 equal quadrants and 1 image per quadrant (4 total) was taken per colony. Using AxioVision, if brightness or contrast was amplified, it was done so uniformly across the entire image and to the negative control images to ensure no false positives or negatives were being created. Images were exported as TIFF files and analyzed using ImageJ software. Marker positive cells were assessed by manual counting based on presence or absence of fluorescent signal, and not fluorescent intensity. *In vivo* tissue sections were imaged using an Olympus FV1000 confocal microscope with FluoView software. Tissue was imaged with 1 μm stacks with ∼15-20 stacks per tissue section and z-stacks were visualized with maximum intensity projections.

### RNA extraction and qPCR

RNA from stem or progenitor cell colonies, or differentiated cells was extracted using the Norgen single cell RNA purification kit (Norgen, 51800), and details for extraction protocol can be found in the kit user manual. Purified RNA was quantified using a NanoDrop and stored at −80°C. For cDNA synthesis, approximately equals amounts of input RNA was used for each sample. cDNA was synthesized using the SuperScript IV VILO kit (Thermo, 11756050) and instructions can be found in the kit user manual. Due to using probes that span exons, DNase treatment was omitted. cDNA was diluted 1:10 in DNase-free RNase-free H2O (Thermo, 10977015) for use in qPCR reactions. The protocol for TaqMan Universal PCR master mix (Thermo, 4304437) and TaqMan probes was used for qPCR reactions on a QuantStudio 6 Flex real-time PCR machine. The following Thermo TaqMan probes with FAM-MGB dyes were used: Gapdh (Mm99999915_g1), Actb (Mm02619580_g1), Vsx2 (Mm00432549_m1), Rho (Mm01184405_m1), Opn1sw (Mm00432058_m1), Crx (Mm00483995_m1), Sox15 (Mm00839542_g1), Casz1 (Mm01258841_m1), and Ikzf1 (Mm01187877_m1). ΛΛCt method was used for relative gene expression and Gapdh and Actb were used as endogenous control genes in these calculations.

### Bulk RNA sequencing

E14 RPCs were cultured in colony forming sphere assays and then differentiated in rod or cone photoreceptor conditions as described in “Primary stem and progenitor cell culture and differentiation”. Each timepoint and culture condition had 3 independent biological replicates. For COCO or T + RA primed spheres, E14 RPCs were cultured at clonal density in SFM + bFGF + heparin with COCO or T + RA, and non-primed spheres were cultured in regular SFM + bFGF + heparin colony forming conditions. RNA was extracted using the Takara pico input mammalian RNA extraction kit (634411) and each sample had high quality RNA with a RIN of 9 or above. Library prep was done with Takara low input RNA kit (634938). Sequencing analysis was done using a published open-source software package (Pertea et al., 2016). Briefly, HiSAT2 was used to align reads to GRCm38 with Ensembl GTF, transcripts were assembled using StringTie, raw read count assessed with HTseq, and differential expression analysis was done using Ballgown. For lists of differentially expressed genes, only known protein coding genes with a q value (minimum false discovery rate) of <0.05 were considered. Gene expression values were used for Principal Component Analysis by “procomp” in base R. The top 3 PCs were used for 3D plots by “plot3d” as part of the “rgl” package in R with designated groups labeled by color.

### Statistical Analysis

Graphs were created and statistical tests were performed using Prism 9 software. For all graphs error bars represent mean ± S.E.M. Replicate number, statistical tests used, and results of those tests are present in each figure legend.

## Data Availability

Bulk RNA sequencing data has been deposited in the NCBI Gene Expression Omnibus (GEO) under the accession number GSE283880.

## Lead Contact

Further information and requests for resources and reagents should be directed to and will be fulfilled by the lead contact, Justin Belair-Hickey.

## Competing Interests

The authors declare no competing or financial interests.

## Acknowledgments

We thank all members of the van der Kooy laboratory for their helpful discussion of concepts, data, and comments on the manuscript. The authors thank Neil Winegarden and Gurbaksh Basi from the Princess Margaret Genomic Centre for their services in sequencing RNA and Zhibin Lu from the UHN Bioinformatics and HPC Core for services in initial bioinformatic analysis of sequencing data.

## Author Contributions

J.J.B.H designed and carried out experiments, analyzed data, and wrote the paper. S.K designed and carried out experiments and analyzed data. B.L.K.C provided technical assistance with dissections. B.B designed experiments and analyzed data. J.C.L compiled and analyzed sequencing data. G.B designed experiments and analyzed sequencing data. D.v.d.K designed experiments, analyzed data, and wrote the paper.

## Funding

Supported by grants from the Krembil Foundation and the Canadian Institutes of Health Research (D.v.d.K).

---

**Fig. S1.**
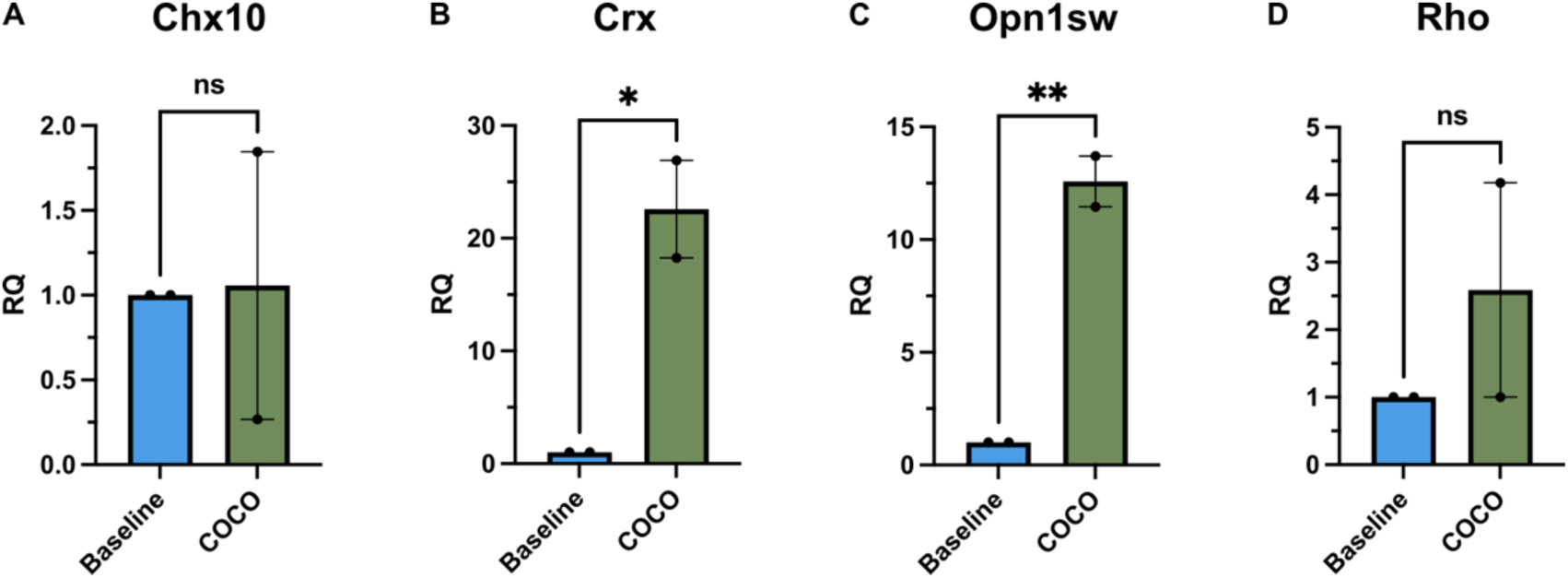
COCO promotes cone differentiation in 3D organoid cultures. **(A-D)** qPCR on day 14 mouse retinal organoids derived from mESCs. Organoids were grown in the presence or absence of COCO and data is shown as fold change (RQ) compared to baseline organoid medium (without COCO). *Gapdh* and *Actb* were used as endogenous control genes. *N* = 2 independent experiments. Two-tailed unpaired Student’s t-test; **A**, *p* = 0.9490; **B**, *\*p* = 0.0379; **C**, *\*\*p* = 0.0094; **D**, *p* = 0.4226. Error bars represent mean ± SEM. For all micrographs main image scale bar is 100 μm and magnified scale bar is 25 μm.

**Fig. S2.**
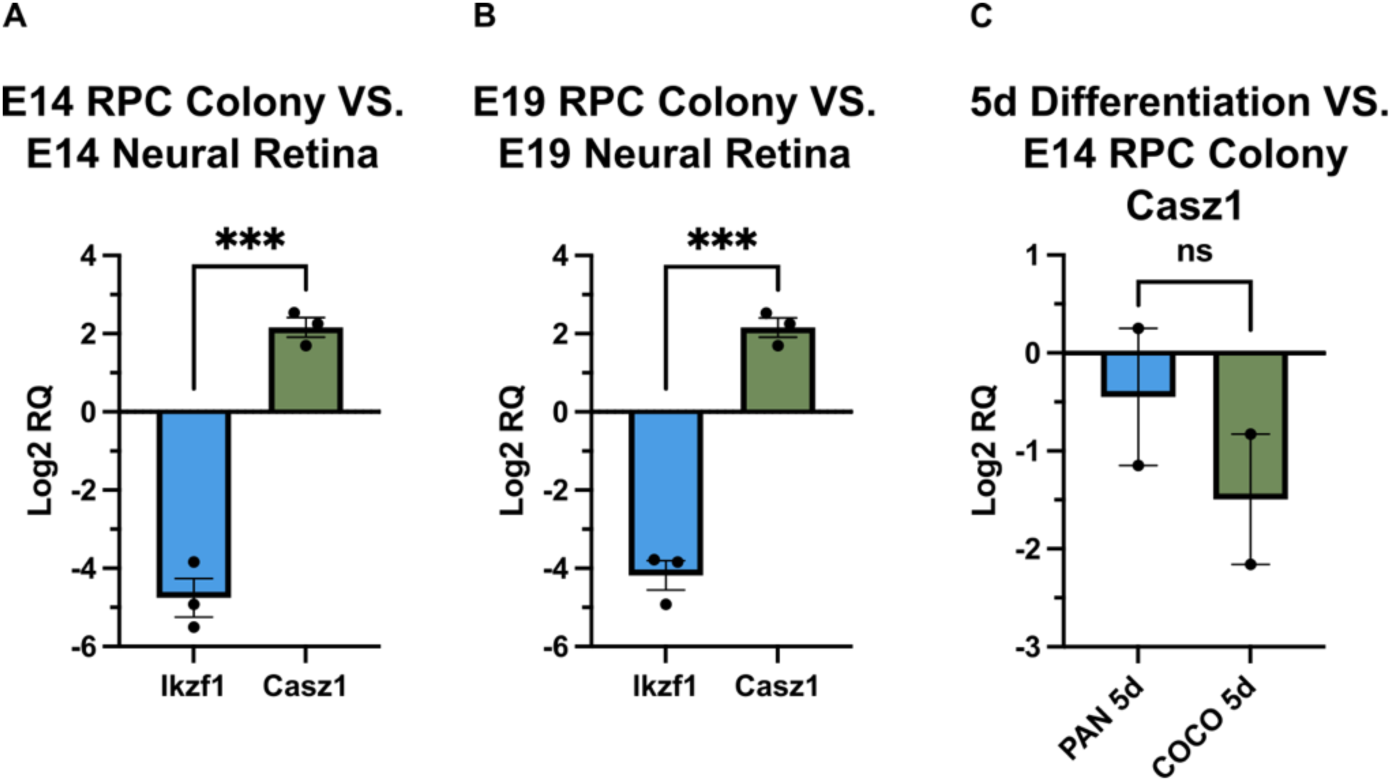
Cell culture conditions alter temporal identity markers in RPCs. Single embryonic RPCs were grown into free-floating spherical colonies for 6 days in culture. qPCR was then used to compare expression of *Ikzf1* (early retinal temporal identity factor) and *Casz1* (late retinal temporal identity factor) relative to whole neural retina at the time in which RPCs were derived. *Gapdh* and *Actb* were used as endogenous control genes. **(A)** Log2 fold change (RQ) of E14 RPC spheres relative to whole E14 neural retina. *Izkf1*, *n* = 3; *Casz1*, *n* = 3; two-tailed unpaired Student’s t-test, *\*\*\*p* = 0.0002. **(B)** Log2 RQ of E19 RPC spheres relative to whole E19 neural retina. *Izkf1*, *n* = 3; *Casz1*, *n* = 3; two-tailed unpaired Student’s t-test, *\*\*\*p* = 0.0001. **(C)** Individual RPC colonies derived from E14 neural retina were differentiated in PAN or COCO conditions for 5 days. *Izkf1*, *n* = 2; *Casz1*, *n* = 2; two-tailed unpaired Student’s t-test, ns, *p* = 0.3923. Error bars represent mean ± SEM.

**Fig. S3.**
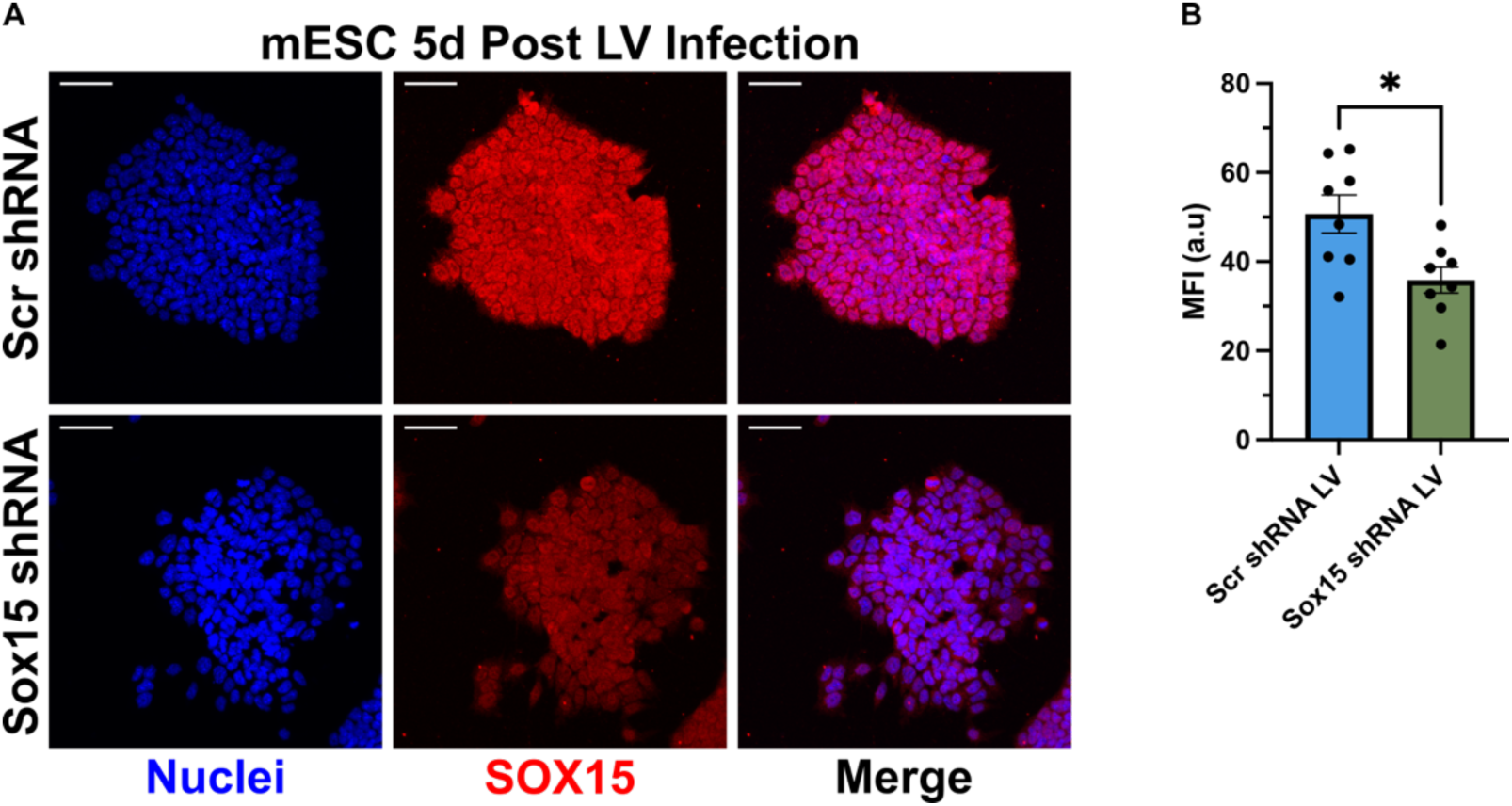
Lentiviral shRNA can significantly reduce SOX15 protein levels. **(A)** Mouse embryonic stem cell colonies (mESCs) were transduced with lentivirus (LV) containing scrambled control (Scr) or Sox15 shRNA constructs and imaged 5 days post transduction. Representative immunofluorescent micrographs show SOX15 protein in a single mESC colony. For all micrographs scale bar is 100 μm. **(B)** Quantification of mean fluorescence intensity (MFI) in individual mESC colonies (described in **A**) expressed in arbitrary units (a.u). Scr shRNA LV, *n* = 8 colonies; Sox15 shRNA LV, *n* = 8 colonies; two-tailed unpaired Student’s t-test, *\*p* = 0.0122. Error bars represent mean ± SEM.

**Fig. S4.**
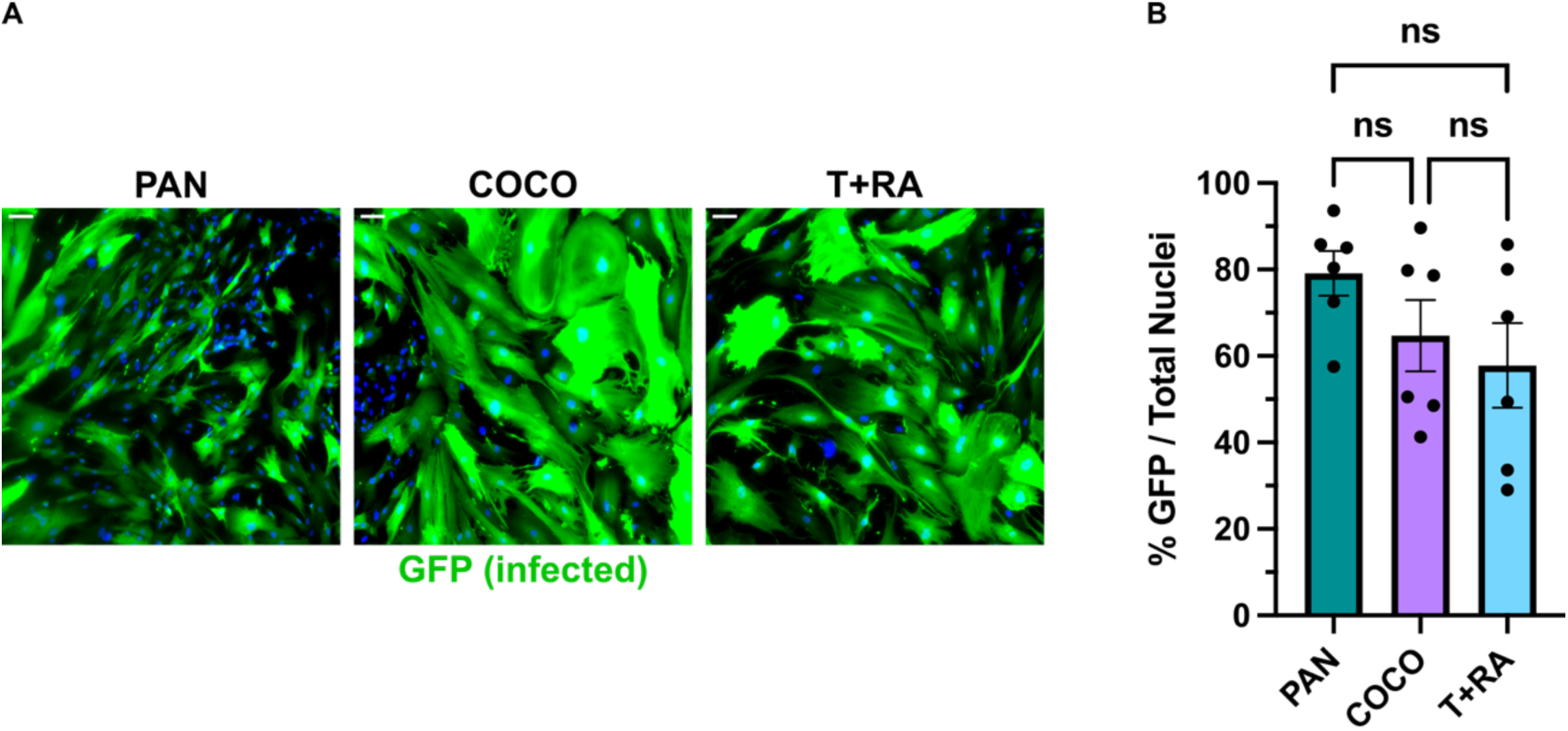
Differentiation condition does not affect transduction efficiency. **(A)** Representative immunofluorescent micrographs of aRSC clones transduced with an EGFP LV and differentiated in PAN, COCO, or T + RA. **(B)** Percentage of total cells in each clone in each differentiation condition that are GFP-positive when transduced with an EGFP LV. PAN, *n* = 6 clones; COCO, *n* = 6 clones; T + RA, *n* = 6 clones; one-way ANOVA with Tukey’s multiple comparisons test, ns, *p* >0.05. Error bars represent mean ± SEM. For all micrographs, scale bar is 100 μm.

**Fig. S5.**
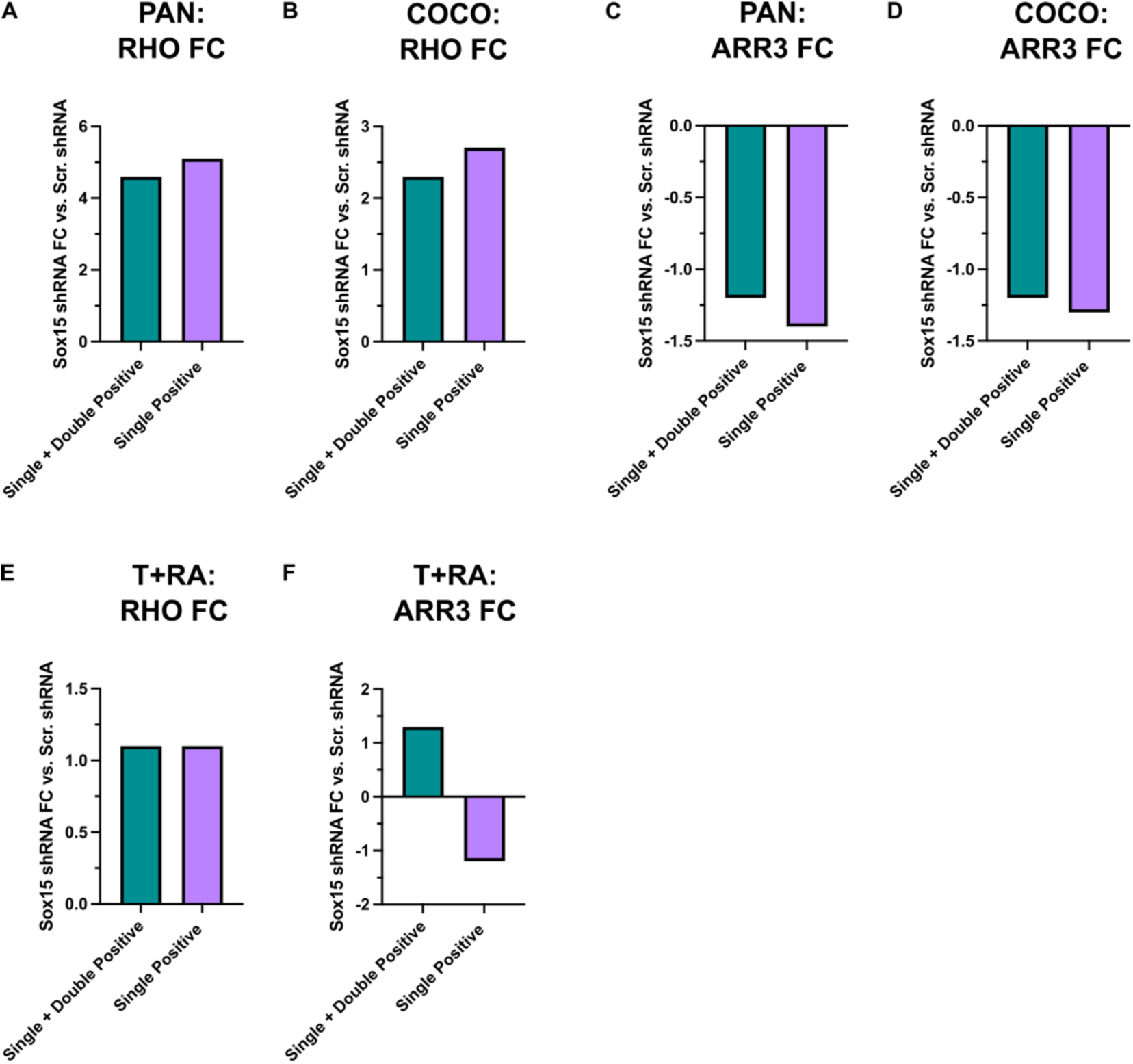
Photoreceptor fold-change with *Sox15* KD. F**(A-F)** old change (FC) in the mean cone (ARR3) and rod (RHO) staining percentages in all three differentiation conditions from Fig. 8. Data represents the fold change with *Sox15* shRNA KD compared to Scr. shRNA. “Single + Double Positive” groups are the mean percentage of total nuclei quantifications in Fig. 8A-I and do not consider cells that are double positive for ARR3 and RHO. “Single Positive” groups have subtracted the mean double positive cells (in Fig. 8J-N) from the mean rod or cone percentage quantifications in Fig. 8A-I.

## Notes

### Competing Interest Statement

The authors have declared no competing interest.

### Summary of Updates

Larger magnified images in Figure 1 and outlined anatomical regions of the retina in Figures 5 and 6.

